# Adapting protein language models for structure-conditioned design

**DOI:** 10.1101/2024.08.03.606485

**Authors:** Jeffrey A. Ruffolo, Aadyot Bhatnagar, Joel Beazer, Stephen Nayfach, Jordan Russ, Emily Hill, Riffat Hussain, Joseph Gallagher, Ali Madani

## Abstract

Generative models for protein design trained on experimentally determined structures have proven useful for a variety of design tasks. However, such methods are limited by the quantity and diversity of structures used for training, which represent a small, biased fraction of protein space. Here, we describe proseLM, a method for protein sequence design based on adaptation of protein language models to incorporate structural and functional context. We show that proseLM benefits from the scaling trends of underlying language models, and that the addition of non-protein context – nucleic acids, ligands, and ions – improves recovery of native residues during design by 4-5% across model scales. These improvements are most pronounced for residues that directly interface with non-protein context, which are faithfully recovered at rates >70% by the most capable proseLM models. We experimentally validated proseLM by optimizing the editing efficiency of genome editors in human cells, achieving a 50% increase in base editing activity, and by redesigning therapeutic antibodies, resulting in a PD-1 binder with 2.2 nM affinity.

## Introduction

Protein sequence design aims to identify an amino acid sequence that will fold into a desired backbone and carry out a function of interest. The ability to design proteins with novel functions has broad applications in biotechnology and medicine, including the development of therapeutics, vaccines, and industrial enzymes. Physics-based methods, such as Rosetta (1), approach protein design as an optimization problem, searching for sequences that minimize an energy function. However, these methods are limited by the accuracy of the underlying energy function and are computationally expensive in practice. Recently, deep learning approaches, which learn a mapping from structure to sequence, have emerged as an alternative to physics-based methods (2–4). Generative models for protein design, such as ProteinMPNN (3), have proven successful across a variety of tasks, including design of protein binders (3), assemblies (5), diversified enzymes (6), and conformational switches (7). Despite their success, protein design models trained solely on experimentally determined structures are ultimately limited by the quantity and diversity of their training data. While the Protein Data Bank (PDB) (8) contains over 200,000 protein structures, this number is dwarfed by the number of sequences identified through genomic and metagenomic sequencing efforts. To overcome this disparity, predictions from AlphaFold2 (9) have been used to supplement the structures used for training (4). This approach has been shown to improve the performance of protein design models (4), but is still limited by the number of structures that can be predicted and the accuracy of the predictions. Protein language models present an alternative means of modeling protein sequence-function relationships.

These models learn directly from sequences through self-supervised training objectives, such as masked residue prediction and next-residue prediction. With increasing numbers of parameters, protein language models have been shown to capture properties including structure (10, 11) and function (12, 13). Design with protein language models is largely a challenge of steering generation towards a desired region of sequence space. Steering of generated sequences is typically achieved through fine-tuning on curated datasets of family- or function-specific proteins. This approach has proven successful for a variety of enzyme families, including lysozymes (14), carbonic anhydrases, lactate dehydrogenases (15), and complex genome editing systems (16). However, the dependence on natural examples for fine-tuning currently limits the scope of design with protein language models to known protein families and functions. Further, it does not offer a straightforward means of incorporating precise atomistic information into the design process, instead relying on the model to implicitly learn these constraints from the fine-tuning data.

In this work, we describe proseLM (protein structure-encoded language model), a method for steering protein language models for design tasks by providing explicit structural and functional information. ProseLM leverages parameter-efficient adaptation of protein language models to incorporate structural and functional information, including the backbone-of-interest and nearby context (proteins, nucleic acids, ligands, ions, etc.). ProseLM effectively incorporates this context while benefiting from the scaling trends of the underlying language models, enabling high rates of native sequence recovery. We experimentally validate proseLM by designing functionally improved genome editors and therapeutic antibodies.

## Results

### Protein structure-encoded language model

Our method for protein sequence design, proseLM, leverages pre-trained protein language models for structure-conditioned design. We take the ProGen2 language models (13) as a foundation and propose a conditional adapter for parameter-efficient incorporation of structural and functional context.

Several methods have been proposed for parameter-efficient fine-tuning of large language models (17–19), enabling specialization of models by training a small fraction of the total parameters. In this work, we build on adapters, which introduce a set of bottlenecked operations that modify the outputs of each model layer. For proseLM, we follow Houlsby et al (18) and introduce adapter layers that update the outputs of the simultaneous attention and feed-forward operations of each ProGen2 layer (Figure 1A). With sufficiently reduced bottleneck dimension, the number of parameters in these adapter layers are miniscule with respect to the pre-trained model. To inject structural information into the ProGen2 language models, we propose an adapter architecture that conditions in the low-rank representation using a multi-layer perceptron (MLP). In this way, the adapter layers maintain parameter efficiency while incorporating structural context throughout the depth of the language model.

**Figure 1.**
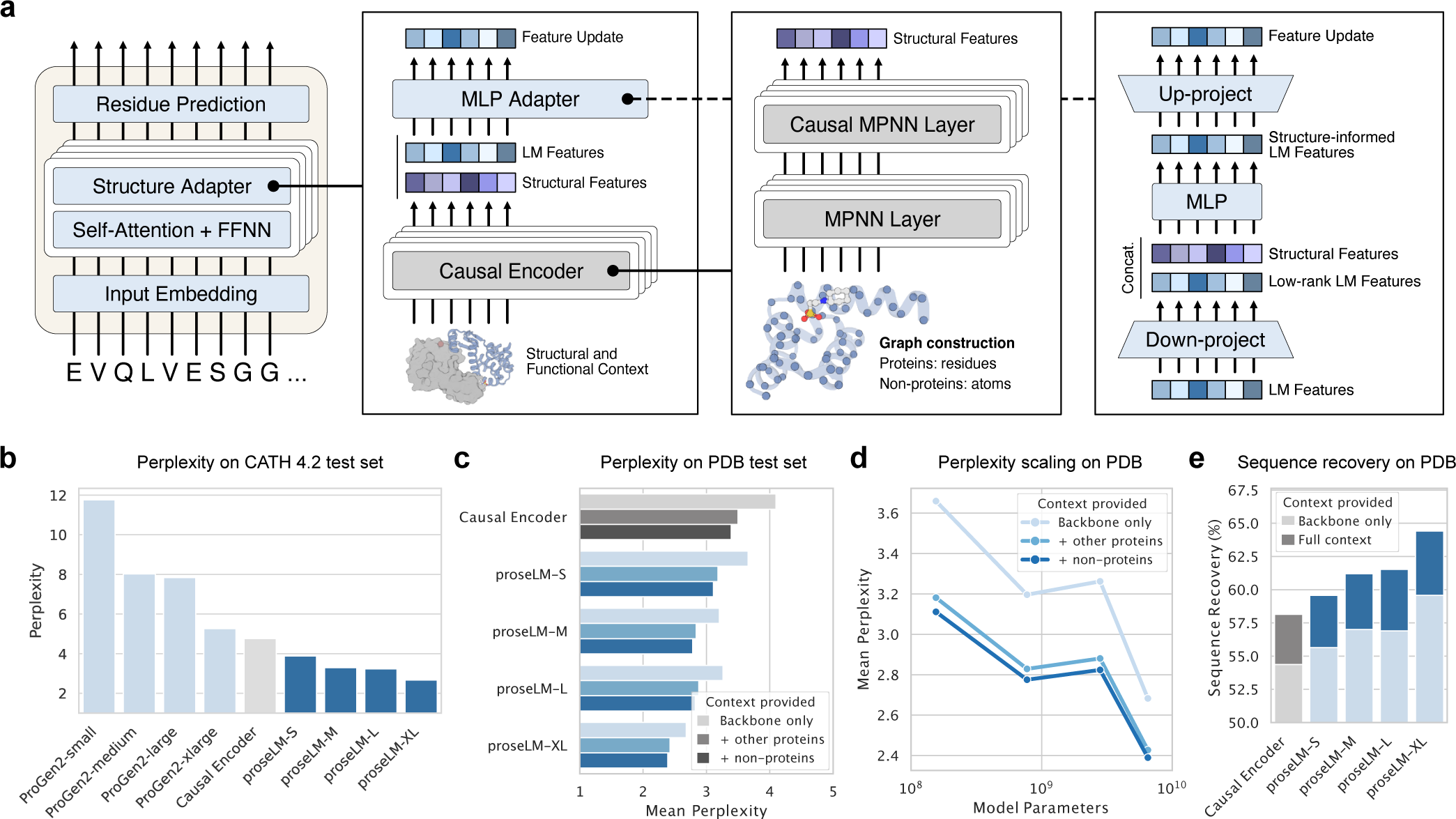
Design of protein sequences for diverse backbones. (a) Diagram of proseLM architecture. Structural adapter layers are placed after each layer of the pre-trained language model. In each adapter layer, structural context from the causal encoder is combined with language model embeddings to condition sequence generation. (b) Perplexity of ProGen2 pre-trained language models and single-chain proseLM models on the CATH 4.2 test set collected by Ingraham et al. (2). (c) Perplexity of proseLM models on the clustered PDB dataset collected by Dauparas et al. (3). Perplexity is reported as the mean of cluster-averaged values. For all models, perplexity decreases as additional structural and functional context is provided. (d) Perplexity with respect to total model parameters for proseLM models trained on the PDB with increasing levels of structural and functional context provided. (e) Native sequence recovery for fully designed sequences from proseLM models. Sequence recovery is reported as the median of cluster-averaged values. For all models, native sequence recovery increases when structural and functional context is provided.

Structural context for conditioning is obtained from a pre-trained causal encoder, which is trained in a similar fashion to prior encoder-decoder protein design models. The causal encoder architecture consists of alternating message-passing (MPNN) and invariant-point message-passing (IPMP) (20) encoder layers (eight total) followed causally masked MPNN and IPMP layers (four total). IPMP layers modify standard MPNN layers by adding frame-based inter-residue features, and have been proposed as a drop-in replacement for protein side-chain modeling and sequence design (20). Following ProteinMPNN, the causal encoder is trained to decode randomly permuted sequences given a fixed backbone (3). Different from the natural N-to-C fixed decoding order, this formulation enables conditioning on later residues for design tasks where some of the sequence should remain constant. We additionally train the causal encoder with masked structural spans, following the span sampling scheme used for ESM-IF1 (4). When trained on the CATH 4.2 dataset (2, 21), the causal encoder achieves a median sequence recovery of 47.24% and perplexity of 4.76 (Table S1) on the test set. Taken together, this performance is similar to ProteinMPNN (trained without noise) (3), which achieves a median sequence recovery of 45.96% and perplexity of 4.61 on the same proteins (22).

To implement proseLM, we combined the pre-trained causal encoder with pre-trained ProGen2 language models (13) using parameter-efficient adapter layers. Standard adapter layers are typically composed of a feed-forward network with a bottleneck dimension, expressed as (18):

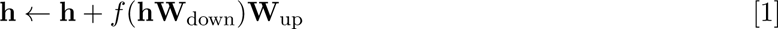

where *h* is the language model embeddings, *f* is a non-linear activation function (typically ReLU), and *W*_down_ and *W*_up_ are weight matrices for the down- and up-projections, respectively. For proseLM, we propose a conditional adapter architecture that incorporates structural context into the language model embeddings using an MLP. The conditional adapter layers are expressed as:

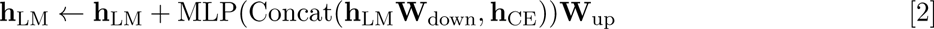

where *h*_LM_ and *h*_CE_ are the language model and causal encoder embeddings, respectively. The causal encoder embeddings are taken from the last decoder layer, just prior to amino acid prediction. By concatenating the causal encoder embeddings with the low-rank language model embeddings, the conditional adapter maintains the parameter-efficiency of standard adapters while incorporating conditioning information. Conditional adapter layers are placed after each of the simultaneous attention and feed-forward layers of ProGen2, with separately trained weights for each adapter layer. During training, the parameters of the causal encoder and conditional adapter layers are updated, while the parameters of the language model are frozen.

To test the effectiveness of the conditional adapters for structure-conditioned sequence modeling, we compared the perplexity of proseLM models trained on the CATH 4.2 dataset to their constitutive models. For ProGen2 models, perplexity scaled with model size, with ProGen2-xlarge (6.4B parameters) approaching the performance of the structure-aware causal encoder (Figure 1b). For proseLM models, perplexity further improved over the causal encoder, following the same scaling trend as the underlying language models. We next compared the native sequence recovery of proseLM models to the causal encoder. Native sequence recovery is defined as the percentage of designed residues that match the native sequence for a particular structure. Native sequence recovery increased with proseLM model scale, with proseLM-XL achieving a 3.59% higher median recovery rate than the causal encoder (Figure S2a). This increase in sequence discovery was distributed across surface and core residues (Figure S2b) and was most dramatic for larger proteins (Figure S2c).

Training structure-conditioned sequence design methods with Gaussian noise applied to the coordinates has been shown to increase robustness of single-sequence AlphaFold2 predictions, which can serve as a proxy for design quality (3). To assess the impact of training with coordinate noise on proseLM, we trained an additional set of causal encoder and proseLM models with 0.1 Å Gaussian noise added to the backbone coordinates. As reported for ProteinMPNN, we found that all proseLM models trained with coordinate noise achieved higher rates of single-sequence prediction structure prediction success with AlphaFold2 (9) and yielded more confident structures (Figure S3). Interestingly, these improvements were most pronounced for the causal encoder and smaller proseLM models, suggesting that the larger models are more robust to structural noise.

### Functional context constrains design space

Protein function depends on interactions with other molecules, ranging from recognition of specific DNA sequences to coordination of metal ions. While historically these interactions have been modeled through physics-based methods (1), they have largely been neglected in recent protein sequence design models. For proseLM, we sought to incorporate functional context beyond the backbone to include not only protein-protein interactions, but also interactions with arbitrary non-protein molecules (e.g., nucleic acids, ligands, ions, etc.). We started by extending the causal encoder to consider protein complexes, which was achieved by simply expanding the existing protein graph to include multiple chains.

Non-protein context is represented as an atomistic graph, with each atom represented as a node with an associated reference frame. Each atomic frame is constructed using the atom-of-interest and its two nearest neighbors (Figure S1b). Nodes are initialized with the types of each atom encoded as node features, along with the distances between the central atom and its neighbors. To incorporate information from the non-protein context graph into the causal encoder, we introduced an IPMP layer for updating the atomistic features, followed by a cross-graph IPMP layer for updating the protein residue features. The cross-graph IPMP layer is similar to the standard IPMP layer, but uses a different set of weights for protein and atomistic nodes, and does not update the atomistic features. The context-aware causal encoder and corresponding proseLM models were trained on the multi-chain dataset used for ProteinMPNN (3) in the same fashion as the single-chain versions, except the causal encoder parameters were frozen during proseLM training to reduce memory requirements.

We first evaluated proseLM on the subset of the PDB test set that contained protein complexes and interactions with some non-protein entity (Table S3). For each protein chain, we computed the perplexity given increasing levels of context: backbone only, with other protein chains, and with full context. In each scenario, perplexity decreased as more context was provided (Figure 1c). This trend held across model scales, suggesting that even the largest language models benefit from explicit functional information (Figure 1d). We next evaluated the capabilities of proseLM for design on the test set structures that had any protein or non-protein interactions. We computed the median native sequence recovery for designs with proseLM when given the backbone only or full context. Increases in sequence recovery from scaling models and providing functional context were largely additive, with proseLM models achieving a consistent 4-5% increase in sequence recovery with full context (Figure 1e).

The most significant increases in native sequence recovery were for residues within 5 Å of nucleic acids, ligands, and ions (Figure 2a). The causal encoder, which is the most directly comparable to recently developed models (23, 24) for context-aware protein sequence design, achieves sequence recovery (with/without context) of 44.44%/54.90%, 50.00%/64.00%, and 46.15%/66.67% for residues near nucleic acids, ligands, and ions, respectively. These results are similar to those reported for CARBonAra (24) and LigandMPNN (23).

**Figure 2.**
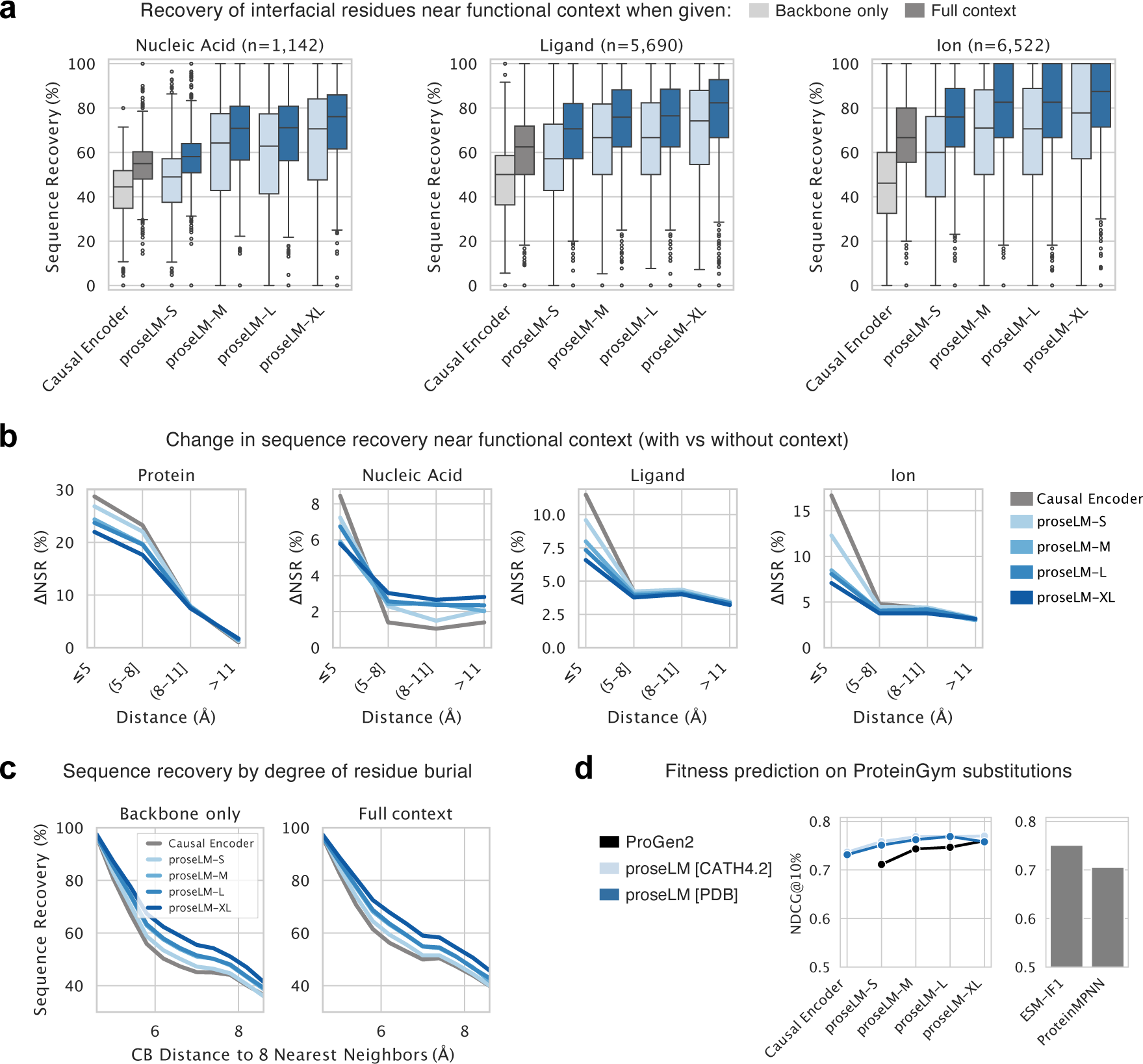
Modeling protein functional context and fitness. (a) Median native sequence recovery for residues near nucleic acids, small-molecule ligands, and ions. (b) Change in native sequence recovery with and without structural and functional context as a function of distance from provided context. Smaller models show the largest increases in recovery near the provided context, while all models converge towards lower overall increases in recovery as distance increases. (c) Sequence recovery as a function of residue burial for protein complex targets. When designed with backbone only (single chain) or with protein context, recovery is largely determined by degree of burial. (d) Evaluation of fitness prediction, measured as normalized discounted cumulative gain, for proseLM models. Overall performance is averaged over landscape categories: stability, binding, activity, expression, and organismal fitness.

With the addition of language model components, proseLM models achieve further increases in native sequence recovery in the vicinity of functional context (Table S3). For ligand and ion interactions, these increases were mostly localized to residues within 5 Å of the provided context (Figure 2b). For nucleic acid interactions, smaller models showed the largest increases in sequence recovery near the provided context, while larger models showed more consistent improvements at greater distances. This suggests that while smaller models benefit from the direct influence of nucleic acid context for local sequence recovery, larger models may be leveraging evolutionary information from pre-training to improve sequence recovery at greater distances. For protein-protein interactions, the effects of context were observable up to 11 Å away, largely due to changes in burial of residues across the designed protein chains (Figure 2c).

### Structure improves modeling of protein fitness

Protein language models trained on evolutionarily diverse sequences are strong zero-shot predictors (i.e., without labeled data) of protein function, with performance on some landscapes benefiting from increased model scale (12, 13, 25). To investigate the impact of structural conditioning on language model fitness prediction, we evaluated proseLM models on a set of 201 deep mutational scan (DMS) datasets curated for ProteinGym (25). We limited our analysis to datasets where the protein sequence length was less than 1024 residues, which was the context length used for training ProGen2 models. Log likelihoods were computed for proseLM models trained on the CATH 4.2 (backbone-only) and PDB (full context) datasets using AlphaFold2-predicted (9) structures from ProteinGym. Datasets are coarsely categorized as stability, binding, activity, expression, or organismal fitness, according to the property measured. Following ProteinGym, we report an overall performance as the average of performance over each category to avoid emphasizing imbalances in the number of datasets available for each fitness type. We additionally report performance for two recent methods – ESM-IF1 (4) and ProteinMPNN (3) – to contextualize the performance of proseLM models.

We quantify fitness prediction performance according to the normalized discounted cumulative gain (NDCG) at ten percent (Figure S5) and the Spearman’s rank correlation coefficient (Figure S6). Although Spearman values are more commonly reported for performance across entire fitness landscapes, the NDCG metric, which focuses on a model’s ability to prioritize the highest-fitness sequences, is more aligned with practical protein engineering settings. Overall, proseLM models achieve higher NDCG values than the causal encoder and ProGen2 models (Figure 2d). Interestingly, the largest improvements in NDCG are for smaller language models, while larger proseLM models often performed comparable to the underlying ProGen2 models. This suggests that larger models gain less new information from structural context, consistent with prior work showing that larger protein language models build increasingly accurate internal representations of structure (11).

For stability landscapes, we observed opposing trends between structure-conditioned and standard language models. The causal encoder was a better predictor of high-stability proteins, while proseLM models showed degraded performance with increased scale. For binding-related fitness landscapes, proseLM models outperformed the causal encoder and underlying ProGen2 models. Interestingly, the backbone-only proseLM models trained on the CATH 4.2 dataset consistently outperformed those trained on the full PDB with additional context. This discrepancy highlights a tradeoff between models that learn to implicitly model functional context and those that explicitly incorporate given context. While the proseLM models trained on the PDB are able to effectively leverage binding partners for prediction, they appear less capable of inferring binding from protein structure alone.

### Experimental validation of functional protein design

Given the compelling performance of proseLM on *in silico* protein design and fitness prediction tasks, we next sought to test its practical utility for functional protein design. Towards this goal, we focused on structure- and function-guided optimization of two broadly relevant protein systems: genome editors and antibodies.

#### Genome editor design

Genome editing technologies repurposed from bacterial anti-phage defense systems have revolutionized life science research and are being actively developed for agricultural and therapeutic applications. In particular, the Cas9 protein from *Streptococcus pyogenes* (SpCas9) has formed the foundation of several downstream editing technologies, including the targeted editing of base pairs in the genome (26). As an RNA-guided endonuclease, the functional activity of SpCas9 is highly dependent on interactions with its guide RNA and the target DNA. We reasoned that the incorporation of structural and functional context into protein language models could enable the local optimization of SpCas9 for increased activity in human cells.

To design variants of SpCas9 with proseLM while maintaining functional activity, we utilized a multi-state conditioning strategy (Figure 3a) that included both the binary and catalytic states of the protein in complex with guide RNA (and DNA for catalytic state). We additionally found it helpful to incorporate evolutionary and functional information in the form of residue-wise conservation patterns from aligned natural sequences and experimental mutation-scanning data, respectively. When combined with multi-state conditioning, this design strategy enabled sampling of low-perplexity sequences within 200 mutations of the wild type sequence (Figure S7). We selected a set of seven designs to test for genome editing in HEK293T cells via co-transfection of the designed proteins and single-guide RNAs (sgRNA) targeting one of three previously characterized target sites. Across all three sites, we observed a wide range of editing efficiencies, with a subset of variants showing activity on-par or higher than SpCas9 (Figure 3b). The most active variant, Variant-2, showed a significant increase in editing compared to wild-type SpCas9 at two of three target sites. Notably, this variant contained considerable mutational load in the REC-1 and HNH domains (Figure 3c), which may facilitate higher on-target editing through reduced specificity and increased nuclease activity.

**Figure 3.**
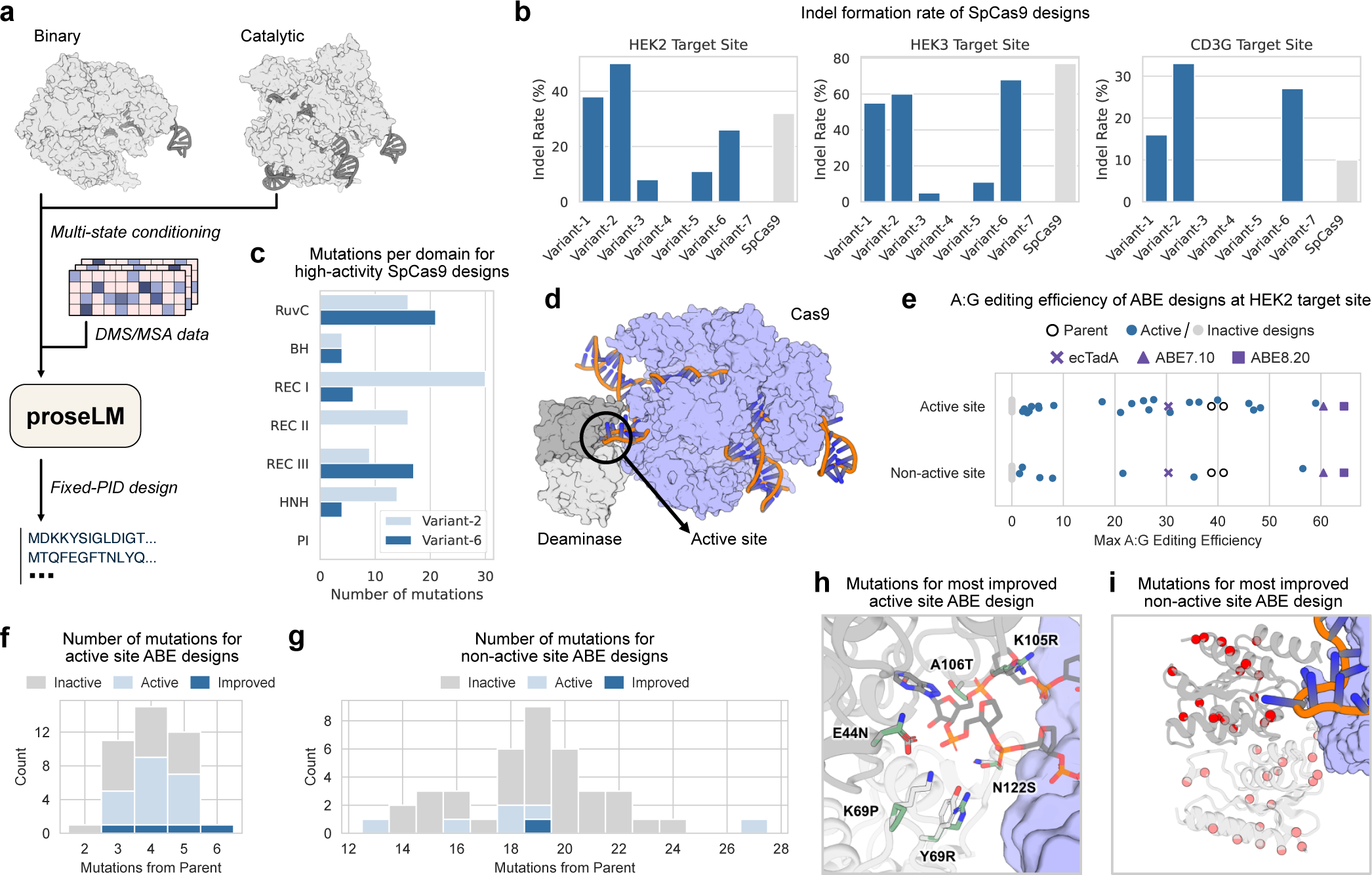
Optimization of genome editors with proseLM. (a) Methodology for design of SpCas9 nuclease variants. Sequences were generated from ProseLM with conditioning on the binary and catalytic states from PDB IDs 4ZT0 and 7Z4J, respectively, and positional residue frequencies from natural sequences (MSA) and experimental data (DMS). The PAM-interacting domain residues were fixed to maintain compatibility with SpCas9 target sites. (b) Editing efficiency of seven designed Cas9 variants across three target sites, with comparisons to parental SpCas9. Five of seven designs showed some activity across at least two guides. (c) Distribution of mutations across SpCas9 domains for two high-activity nuclease designs, with 102 and 59 total mutations. (e) Structure of adenosine base editor (PDB ID 6VPC), with deaminase active site highlighted. (e) Maximum A-to-G editing efficiency for deaminases with design focused on active site or non-active site residues. The parental deaminase activity is indicated by open circles, while active and inactive designs are indicated by blue and gray circles, respectively. The editing efficiencies of three deaminases designed through directed evolution (26, 27) are indicated by purple markers. (f) Number of mutations from parental deaminase for experimentally tested active site designs. Number of designs with observable and improved activity are indicated by light and dark blue bars, respectively. (g) Number of mutations from parental deaminase for experimentally tested non-active site designs. Number of designs with observable and improved activity are indicated by light and dark blue bars, respectively. (h) Structural model of active site six mutations that resulted in highest A:G editing efficiency. Parental residues and mutations are shown in gray and green, respectively. (i) Positions of 19 mutations at non-active site positions that resulted in highest A:G editing efficiency.

We next considered the design of base editors, which are fusions of a deaminase domain to a Cas9 nickase scaffold (26) (Figure 3d). Adenine base editors enable the targeted conversion of A:T base pairs to G:C base pairs in the genome, and have been used to correct pathogenic mutations in human cells (27). As a testbed for optimization with proseLM, we selected a deaminase domain previously designed using protein language models (16) with editing activity on par with early base editors derived from the natural *E. coli* TadA protein (26). We created a structural model of the base editor functional state by aligning an AlphaFold2 (9) prediction of the dimeric deaminase complex to the previously solved structure of ABE8e (28). Using proseLM, we then focused design on active site residues (within 5 Å of the bound adenine) or non-active site residues (outside 5 Å and not in the deaminase dimer interface). We tested 40 designs from each strategy for A-to-G editing efficiency in HEK293T cells and found that both strategies yielded base editors with nearly 50% higher editing efficiency than the parental deaminase (Figure 3e). Among the active site designs, improvements in editing efficiency were achieved with as few as three mutations, while the best design featured six mutations (Figure 3f). For non-active site designs, which ranged from 13 to 27 mutations, fewer designs retained editing efficiency and only one showed improvement over the parental sequence (Figure 3g). In Figure 3h, the set of six active-site mutations yielding a 50% relative improvement in editing efficiency are depicted on the predicted structure of the parental deaminase. While it is difficult to ascertain the precise contribution of each mutation, the design contains several non-conservative mutations, representing a significant reworking of functionally important residues. Meanwhile, the non-active site design with the highest editing efficiency contained 19 mutations distributed across the surface and core of the deaminase (Figure 3i), which may facilitate increased editing through stabilization of the domain.

#### Therapeutic antibody design

Antibodies are a class of immune proteins that have been developed for a wide range of research and clinical applications. The design of specific and biophysically well-behaved antibodies has been a long-standing challenge, due in large part to the complexity and sensitivity of protein-protein interactions typical of antibody-antigen complexes. Recently, protein language models (including antibody-specific models) have been used for targeted optimization of particular antibody attributes, such as stability (29) or immunogenicity (30). However, prior approaches have primarily focused on sequence-based optimization, ignoring the structural context of antibody binding, and focusing on mutations to the framework region. We reasoned that by explicitly modeling the antibody-antigen interface, proseLM would be well-suited for optimization of the binding affinity of therapeutic antibodies.

Although proseLM models effectively recovered native residues at protein-protein interfaces, neither the causal encoder nor the ProGen2 models were exposed to significant numbers of antibody sequences during training. To address this limitation, we trained proseLM-Ab (based on adaptation of the ProGen2-OAS model) on a set of antibody structures from the Structural Antibody Database (SAbDab) (31). We compared the perplexity of ProGen2 and proseLM models on a held-out set of antibodies and found that proseLM-Ab achieved lower perplexity than proseLM models trained on the PDB, including those with significantly more parameters (Figure 4a). These improvements came despite the underlying ProGen2-OAS model assigning relatively high perplexity to the held-out antibodies, likely due to an under-representation of light chains in its training corpus (13, 32). When provided antigen context, all proseLM models assigned lower perplexity to the antibody sequences (Figure 4b) and typically achieved higher rates of native sequence recovery (Figure 4c). For heavy and light chain framework regions, we observed a scaling trend in sequence recovery, with larger models achieving higher recovery rates (Figure 4d). For CDR regions, we observed a similar trend, although the recovery rates were generally lower than for the framework regions. ProseLM-Ab frequently achieved the highest rates of recovery for CDR loops, but showed the largest improvements in the framework regions, demonstrating the utility of adapting antibody-specific language models trained on diverse antibody sequences.

**Figure 4.**
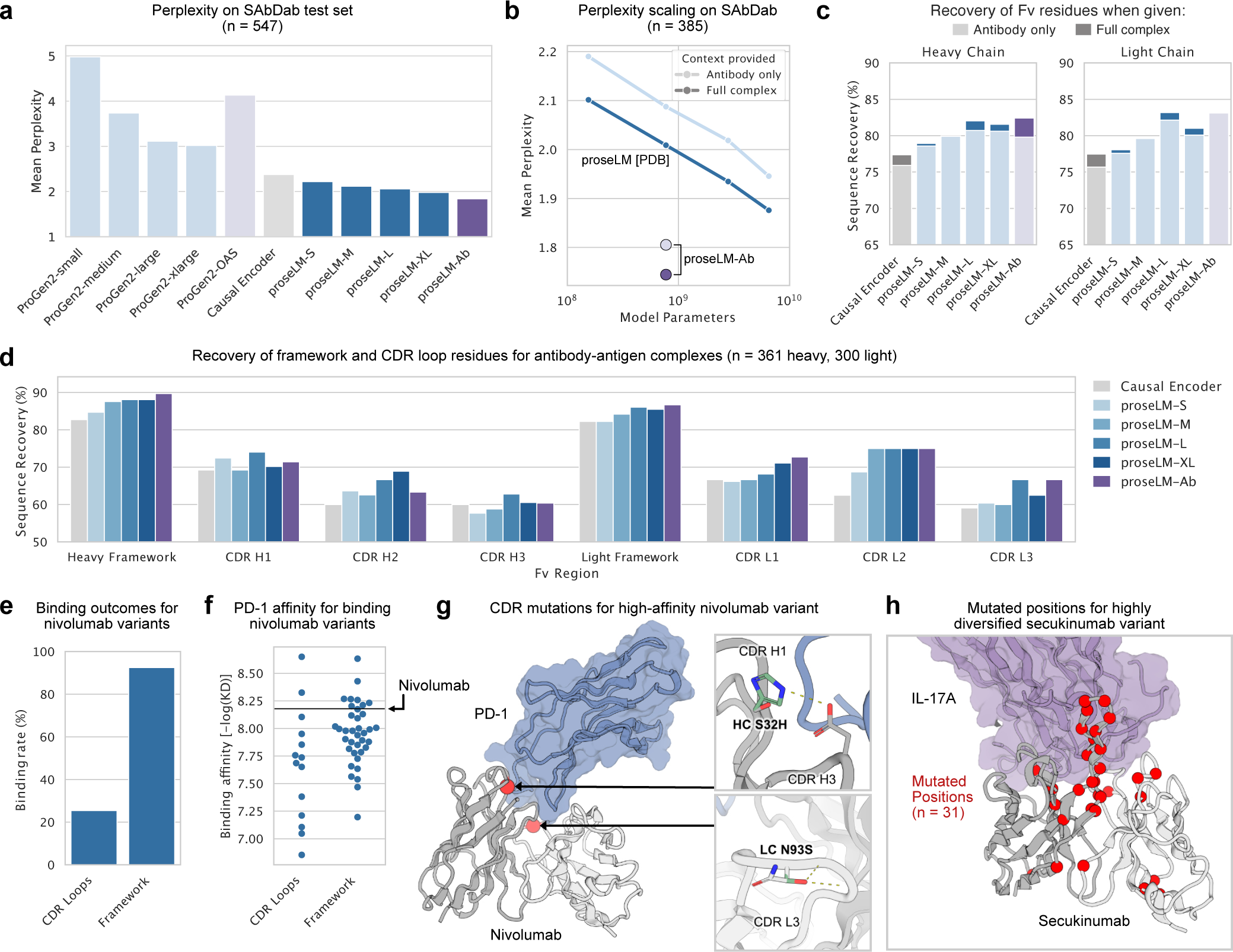
Design of therapeutic antibodies with proseLM. All perplexity and sequence recovery values are reported as the mean and median, respectively, of cluster-averaged values. (a) Perplexity of proseLM models on the antibody dataset collected from SAbDab. For all models, perplexity decreases when antigen context is provided. (b) Perplexity with respect to total model parameters for proseLM models with and without antigen context provided. Blue lines and purple points represent performance of PDB-trained (proseLM [PDB]) and SAbDab-trained (proseLM-Ab) models, respectively. (c) Native sequence recovery for fully designed heavy and light chain variable fragment sequences, with and without antigen context. (d) Sequence recovery for designed heavy and light chain variable fragments by structural region. (e) Percentage of experimentally tested nivolumab variants that retained PD-1 binding for each design strategy. (f) Binding affinity (-log(KD), higher better) for nivolumab variants that retained PD-1 binding. Horizontal line indicates the binding affinity of the parental nivolumab antibody. (g) Structural model of mutations for highest affinity nivolumab variant from CDR-directed optimization strategy. The locations of the two mutations in the Fv are shown as red spheres, with potential novel interactions highlighted. (h) Position of mutations for highly diversified secukinumab variant, with 31 positions mutated across the heavy and light chains.

To test the design capabilities of proseLM models, we first considered affinity optimization of the therapeutic antibody nivolumab, which targets the PD-1 antigen. We focused mutations in either the complementarity-determining regions (CDRs) or the framework regions. For CDR-directed designs, we considered all single and double mutations (excluding mutations to or from cysteine or proline) at positions within 8 Å of the antigen (Figure S9). For framework variants, we redesigned the entire heavy and light chain variable fragments, conditioned on the residues within 8 Å of the antigen (Figure S10). Designs from both strategies were scored using an ensemble of proseLM models and the best 55 CDR-directed and 40 framework-directed variants were selected for experimental characterization. We tested for binding to PD-1 via surface plasmon resonance (SPR) and obtained K*_D_* values for 25.4% of CDR-directed designs and 92.5% of framework-directed designs (Figure 4e). Among these, we observed a wide range of binding affinities, with the most improved variants from each strategy achieving a nearly three-fold increase in binding affinity relative to nivolumab (Figure 4f). The most improved CDR-directed variant contained two mutations near the antigen interface, which likely facilitated tighter binding through improved rigidification of the paratope (Figure 4g), despite one mutation (HC S32H) ablating binding and the other (LC N93S) only moderately improving affinity in isolation. The most improved framework-directed variant contained seven mutations that may have indirectly improved binding through stabilization of the framework (33).

Given the success of targeted optimization of nivolumab binding affinity, we next considered a more aggressive redesign strategy for a more structurally challenging therapeutic antibody. For this task, we selected secukinumab, which binds the IL-17A cytokine through contacts mediated by an extended 18-residue CDR H3 loop. Rather than target specific residues for redesign, we used proseLM to directly redesign the entire heavy and light chain variable fragments of secukinumab (Figure S11). Due to the relative ease of recovering framework residues, the majority of variation was focused within the CDR loops (particularly CDR H3). We tested 96 designs for binding to IL-17A via SPR and found two variants that retained binding (Figure 4h). These variants contained 18 and 31 mutations and bound with 135 nM and 102 nM affinity, respectively. While further optimization of these variants would be necessary to achieve therapeutically relevant binding affinities, these results demonstrate the potential of proseLM for the diversification of structurally challenging therapeutic antibodies.

## Discussion

Recent generative models for structure-based protein design have shown promise for a variety of design tasks (3), but are ultimately limited by the quantity and diversity of structures used for training (4). Protein language models, which learn directly from sequences, have been shown to implicitly capture the structural and functional constraints of proteins (11, 13) and are capable of generating functional proteins (14–16). However, steering protein language models towards a particular fold or function typically requires curation of natural examples for fine-tuning, which may be limited or non-existent for some design tasks. In this work, we presented proseLM, a method for providing explicit structural and functional context to protein language models for structure-conditioned design and showed that the sequence generation capabilities of proseLM benefit from the scaling trends of the underlying language models. Prior work has attempted to circumvent the paucity of experimental structure data by leveraging highly accurate protein structure prediction models, such as AlphaFold2 (9), to generate synthetic training data for protein design models (4). While this approach yielded higher native sequence recovery, the usage of synthetic data may introduce biases that limit the practical utility of the resulting models (34). By contrast, proseLM leverages the wealth of protein sequence data directly, enabling more efficient post hoc incorporation of structural awareness.

In validating proseLM, we selected design tasks where functional context was not only critical, but also where recapitulating the native sequence was not sufficient for success. For optimization of SpCas9, this entailed navigating the local fitness landscape of a protein that is already highly performant. Here, we found that proseLM designs largely retained activity, in some cases improving on-target editing efficiency. Further testing is required to determine the off-target editing profile of these variants, but these results suggest that proseLM can be used to identify functionally relevant mutations in complex protein-nucleic acid systems. For base editor optimization, our starting point was a generated deaminase with activity that already does not exist in nature (deamination of single-stranded DNA). Here, we found that proseLM could be used to directly redesign the active site of the deaminase, yielding variants with significantly increased activity. These results suggest that proseLM can be used in design settings where function deviates from nature, provided a reasonable structural hypothesis. Finally, for antibody design tasks, we began with therapeutic antibodies that are already highly optimized for their intended functions. Against this strong baseline, we found that proseLM could be used to improve binding affinity through CDR-directed or framework-directed optimization, as well as to diversify the binding region of a structurally complex antibody. Overall, these results demonstrate the utility of proseLM for a variety of functional protein design tasks.

Language models have excelled at functional protein design by implicitly modeling structural and functional constraints through fine-tuning on curated examples. As we move towards making these constraints explicit, we anticipate trade offs will emerge in cases where it is difficult to precisely and sufficiently specify the desired functional behavior. In this work, we saw evidence of this tradeoff for fitness prediction, where proseLM models trained with only the protein backbone outperformed those that were explicitly conditioned on functional context during training. This discrepancy was most pronounced for binding datasets, for which fitness is directly tied to an interaction with a specific partner. In light of this, we expect that backbone-only proseLM models will be more useful for design settings where the desired function is challenging to specify structurally, but is consistent with natural evolutionary constraints of the protein family. Meanwhile, for tasks where the desired function is novel or highly specific, we anticipate that explicit conditioning on functional context will be necessary. In the future, models that can incorporate categorical or partial structural conditioning may facilitate the design of proteins with more complex functional requirements.

## Data availability

Protein structure datasets used for training the models described in this work are available from public sources as detailed in the Methods. Designed protein sequences and experimental outcomes are available in the Supplementary Information.

## Code availability

Original code and trained models described in this work will be made available at https://github.com/P rofluent-AI/proseLM-public.

## Author contributions

J.A.R. conceived of the project. A.M. supervised the project. J.A.R. developed the methodology and conducted the investigation. A.B. contributed to the methodology. A.B., J.B., and S.N. contributed to the design and selection of Cas9 proteins. J.A.R. performed design and selection of base editors and antibodies. J.G. designed wet lab experiments. J.R., E.H., R.H., and J.G. performed wet lab characterization of genome editing proteins. S.N. processed experimental genome editing data. J.A.R. wrote the first draft of the manuscript. All authors wrote and/or reviewed the final draft of the manuscript.

## Competing interests

The authors are current or former employees, contractors, or executives of Profluent Bio Inc and may hold shares in Profluent Bio Inc.

## Safety and ethics

Computational protein design carries the dual potential to accelerate development of novel therapeutics and other society-improving molecules, while providing parallel capabilities for nefarious uses, such as engineering of bioweapons. When bolstered by current and future iterations of generative AI, these capabilities are heightened and expected to grow further. The global protein design community has begun to establish appropriate regulations and guidelines towards the continued beneficial development and application of these technologies. In support of these efforts, J.A.R. and A.M. have joined as signatories on a set of community values, guiding principles, and commitments for the responsible development of AI for protein design (https://responsiblebiodesign.ai/). Gene synthesis represents a critical step in the actualization of designed protein sequences. The International Gene Synthesis Consortium (IGSC) unites major gene synthesis providers under a commitment to screen all incoming orders against known pathogens and potentially dangerous sequences. As a concrete step towards safe application of protein design technology, all gene synthesis work in support of the present study was performed with IGSC members. For all protein design projects, we urge researchers to maintain ethical oversight throughout project initiation, experimental characterization, and subsequent deployment phases to ensure safety and avoid unintended harmful outcomes.

## Methods

### Protein structure datasets

Three datasets were assembled for training proseLM models: single-chain proteins, protein complexes with non-protein context, and antibody complexes. For training on diverse single-chain proteins, we used the CATH 4.2 dataset constructed by Ingraham et al. (2), which has been widely adopted as a benchmark for single-chain protein design methods (3, 22, 35). For training on protein complexes with non-protein context, we adapted the dataset used to train multi-chain variants of ProteinMPNN (3). This dataset was originally constructed by clustering the entire PDB – as of August 2, 2021 – at 40% sequence identity and identifying clusters for testing that did not include proteins that co-occurred in biological assemblies with proteins used for training. We extended this dataset by extracting non-protein context within 5 Å of the protein chains. For training on antibody complexes, we collected all antibody structures from SAbDab (31) – as of July 1, 2023 – and performed clustering at 80% identity on the concatenated heavy and light chain variable fragment sequences with MMseqs2 (36). These clusters were used to divide the dataset into training, validation, and test splits, such that roughly 80%, 10%, and 10% of clusters were allocated to each split, respectively.

### Model architecture

#### Structure featurization

We formulate protein structures as nearest-neighbor graphs, with residues as nodes and inter-residue relationships as edges. The graph adjacency matrix is defined by the proximity of residues in sequential and three-dimensional space. For each residue, we first we take the six closest residues along the sequence then the next thirty closest residues according to C*_α_* distance, for a total of 36 neighbors. The nodes are featurized only with a binary indicator of whether the residue has a backbone structure (i.e., N, C*_α_*, and C atoms). The edges are featurized by the inter-residue distances between pairs of backbone atoms (N, C*_α_*, C, O, and virtual C*_β_*) and an embedding indicating the relative position of neighboring residues along the sequence, up to a maximum of 32 positions in either direction. For inter-chain residue pairs, we set the relative positional embedding to a constant value indicating no sequential relationship, but all other features remain the same.

For atomic-level representation of non-protein context, we adopt a similar graph representation, with individual atoms as nodes and inter-atomic relationships as edges. The adjacency matrix for the atomic graph is constructed by selecting the ten nearest atoms in three-dimensional space. Each atom is represented as a node with an associated reference frame formed by the atom-of-interest and its two nearest neighbors. The nodes are initialized with the types of each atom, along with the distances between the primary atom and its neighbors. The edges of the atomic graph are featurized by the inter-atomic distances between these triplets of atoms. Finally, to incorporate information from the atomic graph into the protein graph, we construct a cross-graph adjacency matrix between each protein residue and the nearest thirty atoms (up to 12 Å away) in the atomic graph. The edges for the protein-atomic graph are featurized by the pairwise distances between the backbone atoms of the protein residues and the atom triplets of the atomic graph.

#### Causal encoder architecture

The causal encoder takes as input the protein and atomic context graphs featurized as described above. All discrete node and edge features (binary structural indicator, relative positional embedding, etc.) are one-hot encoded. All distance-based edge features are encoded by sets of sixteen Gaussian radial basis functions (RBFs) equally spaced between 0 and 20 Å. The respective features for nodes and edges are concatenated then processed by two-layer MLPs to bring them to appropriate dimensionality. As in prior work (2, 3), the causal encoder is composed of a series of sequence-agnostic encoder layers followed by a series of causally masked decoder layers that are ultimately used to predict the amino acid sequence (Algorithm 1). Complete hyperparameters for the causal encoder are provided in Table 1.

**Table 1.**
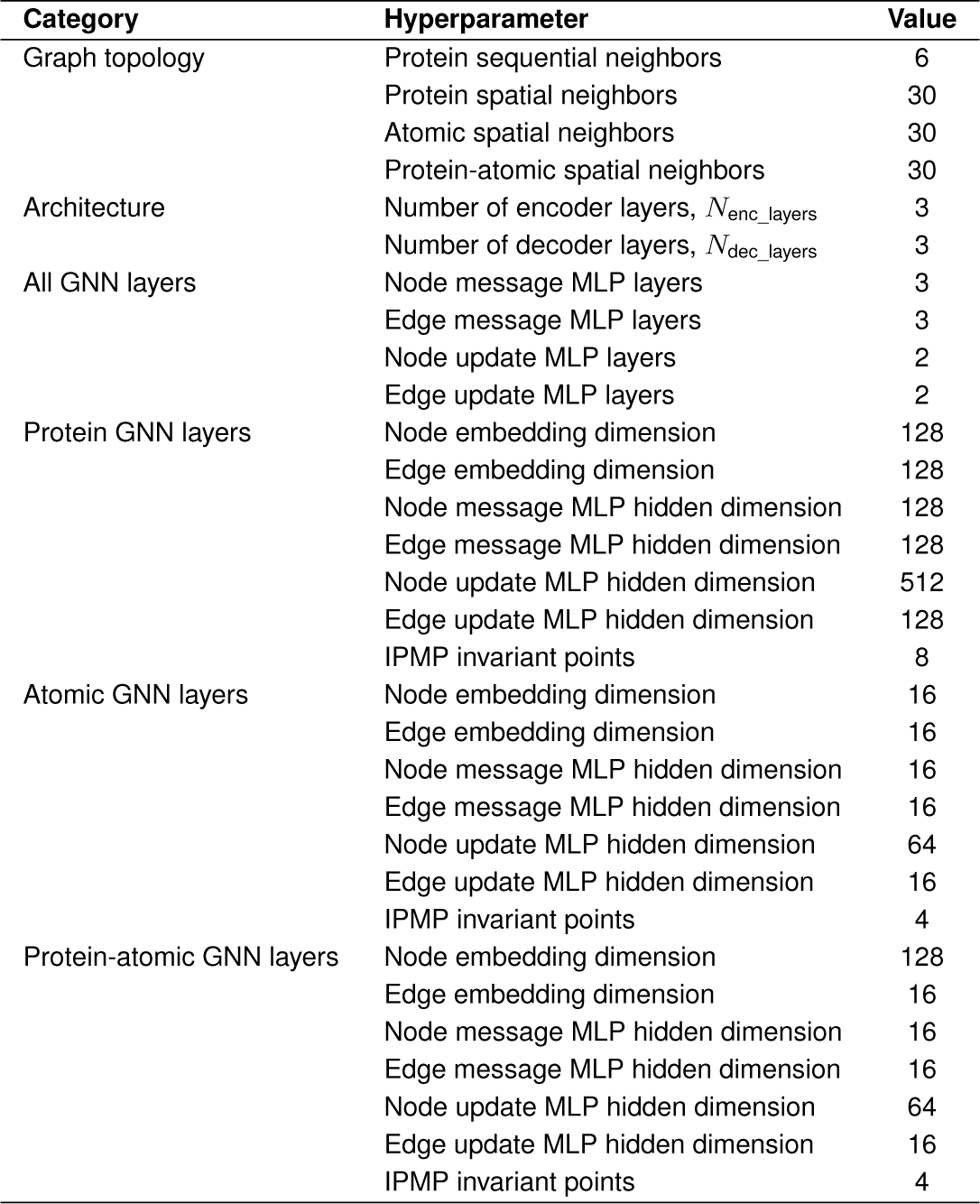
Causal encoder hyperparameters.

The encoder and decoder layers are parameterized by graph neural networks (GNNs), specifically taking the form of message-passing neural network (MPNN) layers (2, 3) and invariant point message-passing (IPMP) layers (20). Our instantiation of MPNN and IPMP layers differ only in the features used to form messages for updating node and edge embeddings. For MPNN layers, messages are formed by concatenating node and edge embeddings for neighbors in the graph topology (Algorithm 2). For IPMP layers, we add to these embeddings the set of five invariant components proposed by Randolph et al. (20) (Algorithm 3). To obtain these components, we predict a set of invariant points in the local frame of each node, then compute the following for each pair of neighboring nodes in the graph (Algorithm 4):

##### Algorithm 1 Causal encoder

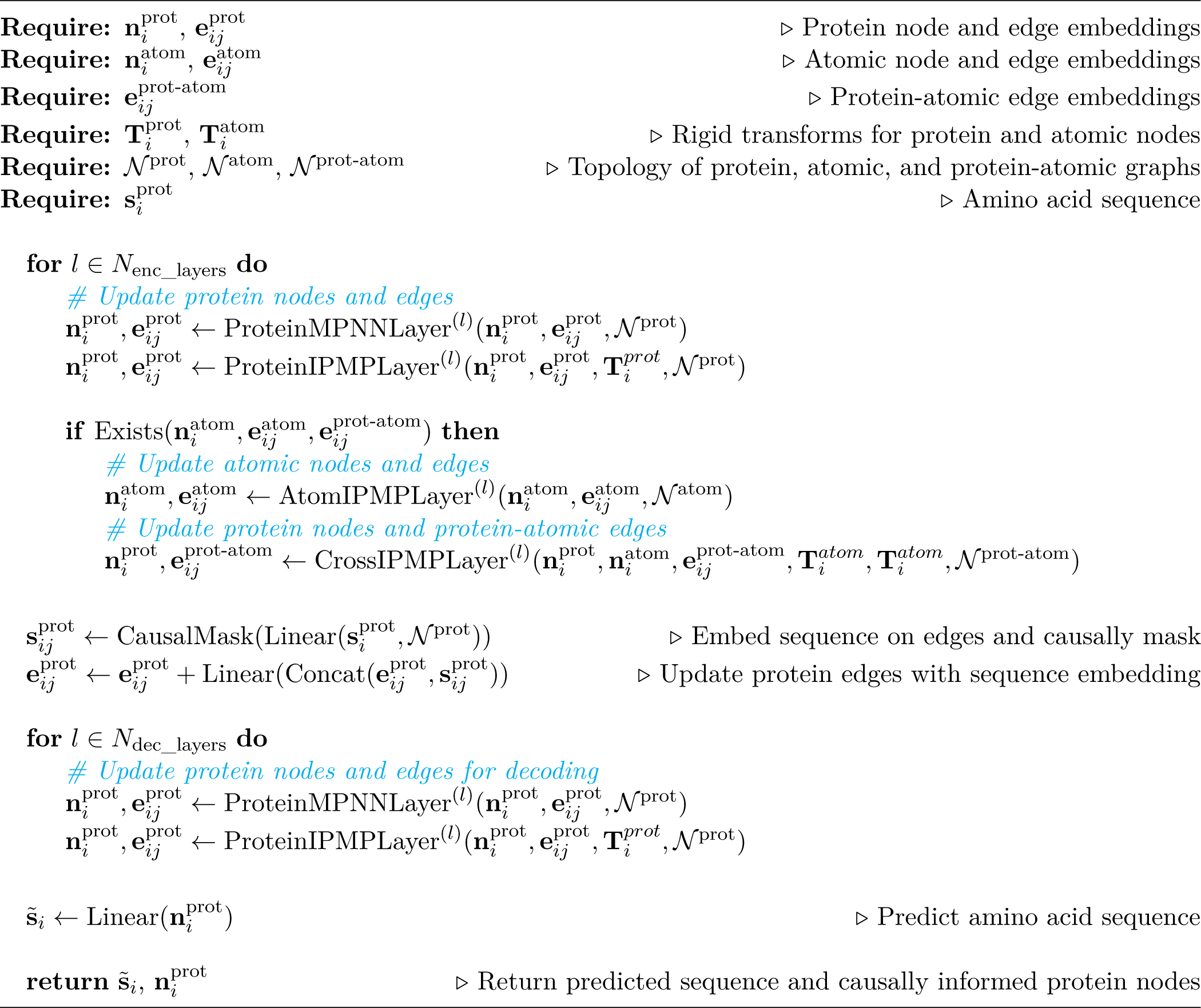

1. Node *i*’s invariant points in node *i*’s local frame
2. Distances from node *i*’s invariants points to the origin of node *i*’s local frame
3. Node *j*’s invariant points in node *i*’s local frame
4. Distances from node *j*’s invariant points to the origin of node *i*’s local frame
5. Distances between node *i*’s invariant points and node *j*’s invariant points in the global frame

After assembling the message features **p***_ij_* for MPNN or IPMP layers, we pass those features through a set of common operations to update node then edge embeddings. For the node embeddings, we compute a set of messages **m***_ij_* for each neighboring node *j ∈ N* using a three-layer MLP and aggregate them by taking the mean across all neighbors. These messages are then added to the original node embeddings and passed through a layer normalization. The updated node embeddings are further processed by a two-layer MLP, with the outputs added back to the previously updated node embeddings with layer normalization. For the edges, we follow a similar procedure but omit the aggregation step and instead directly update the embeddings using the messages.

To enable the decoder layers to predict the amino acid sequence residue-by-residue at inference time, we provide the ground truth (or presently decoded) sequence through causally masked edge embeddings (2). Specifically, for neighboring nodes *i*>*j*, the protein sequence **s**^prot^ is embedded and added to the protein edges. For node and edge updates, we form messages in a causally consistent manner for neighboring nodes *i* and *j* by selectively using encoder node embeddings for node *j* when *i*<*j* and decoder node embeddings when *i*>*j*. The final node embeddings are used to predict the amino acid sequence through a linear layer.

##### Algorithm 2 MPNN layer

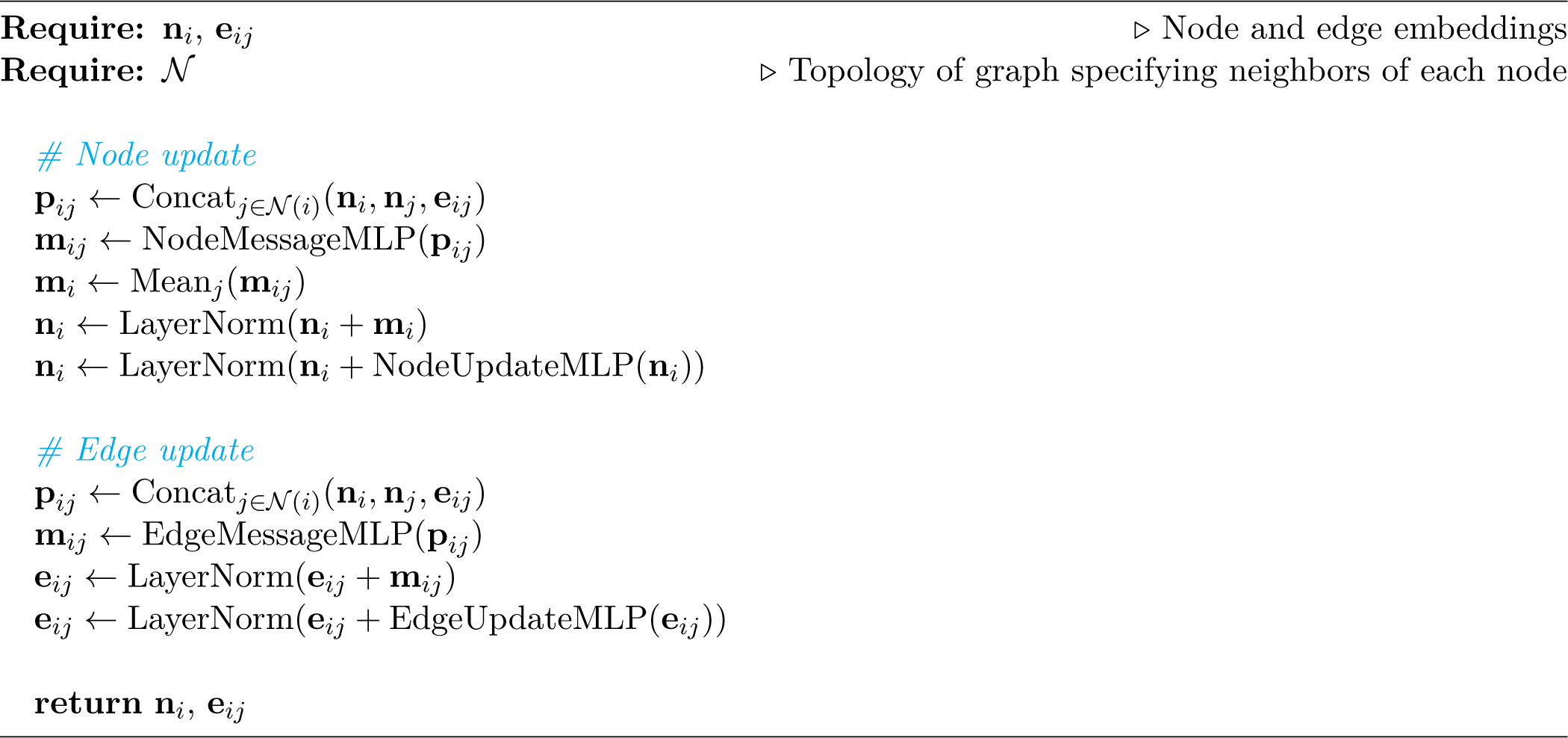

##### Algorithm 3 IPMP layer

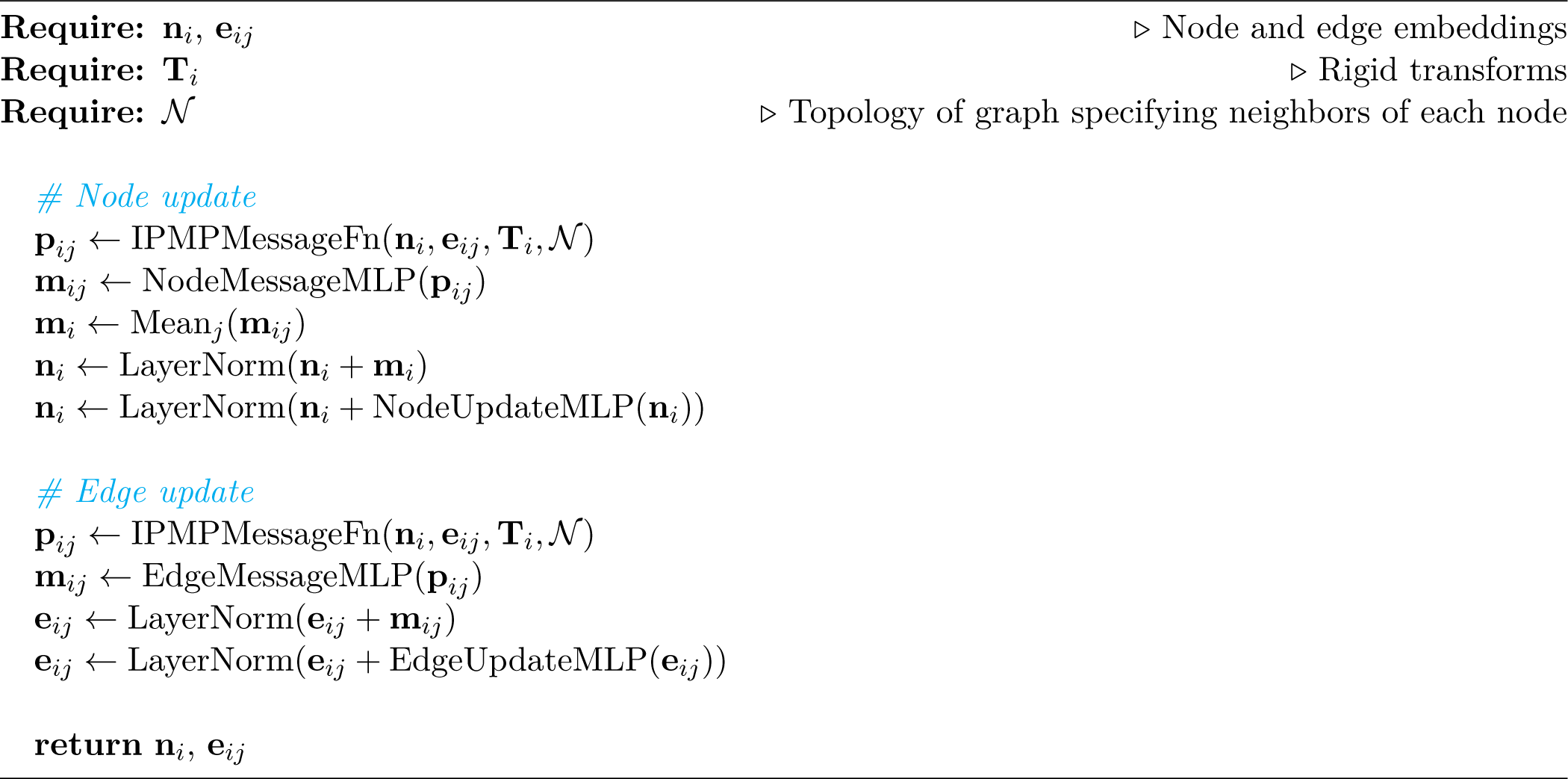

##### Algorithm 4 Invariant point message features

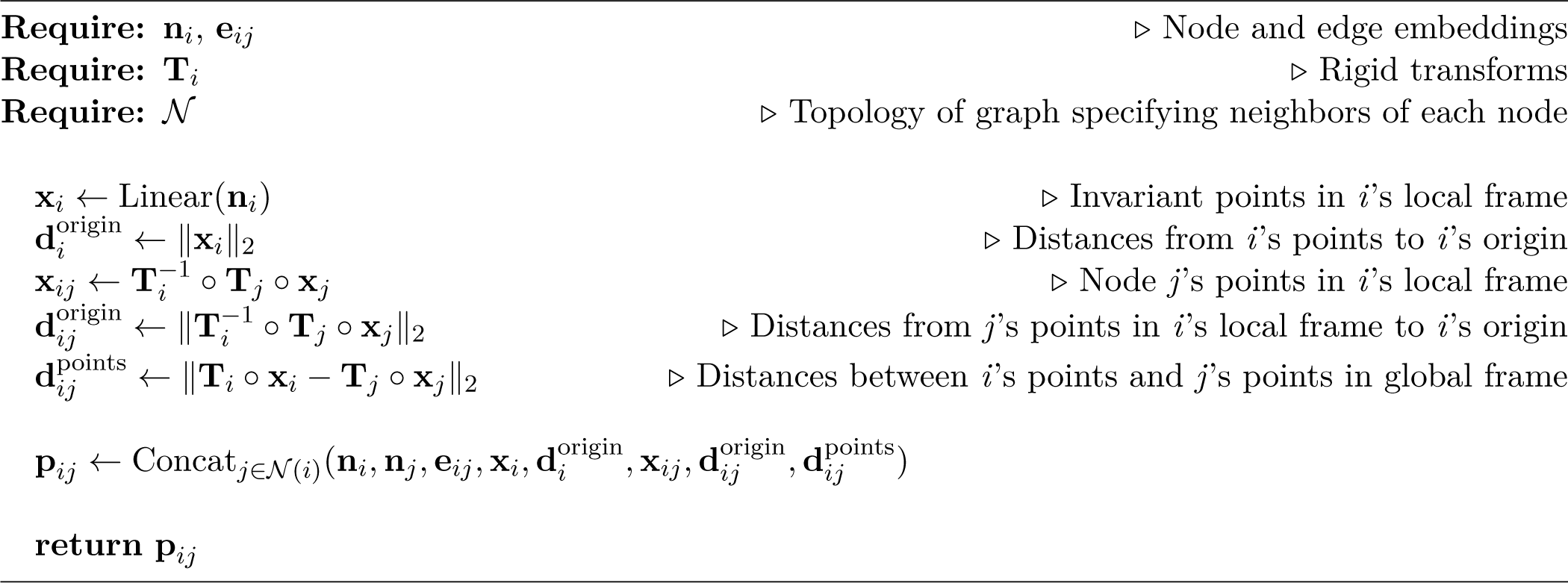

Causal encoder models were trained for 80 epochs on the CATH 4.2 and PDB datasets with an effective batch size of 32 using the Adam optimizer (37). The learning rate was increased linearly over 4,000 warmup steps to a maximum value of 1e-4, then decayed according to an inverse-square-root schedule. An additional set of models were trained in the same manner but with 0.1 Ã Gaussian noise added to the protein backbone coordinates. The final models were chosen according to validation set loss.

#### ProseLM architecture

For proseLM models, we combine a pre-trained causal encoder with a pre-trained ProGen2 protein language model (13) using a set of parameter-efficient conditional adapters (Algorithm 5). The causal encoder is used to encode the protein and atomic structures, as well as the causally masked amino acid sequence, into a set of node embeddings **n**^prot^. These embeddings are then used to condition the outputs of the simultaneous attention and feedforward layers of the language model through a set of MLP adapters (Algorithm 6). The node embeddings are shifted by one position, such that the language model is conditioned on the structural information of the next residue to be predicted, rather than the current residue encoded in the sequence. The adapter layers first down-project the language model embeddings to a reduced dimensionality, then condition on the node embeddings through a two-layer MLP, and finally up-project the conditioned embeddings back to the original dimensionality. Importantly, the linear layer responsible for up-projecting the embeddings is initialized with weights near zero such that the entire adapter layer approximates an identity operation when training begins. The conditioned embeddings are ultimately added back to the original language model embeddings and passed through the pre-trained layer normalization of the language model. Hyperparameters for conditional adapters for each proseLM model are provided in Table 2.

**Table 2.** ProseLM hyperparameters.

| Model | Language model | LM embed. dim. | Reduction factor | Low-rank embed. dim. |
| --- | --- | --- | --- | --- |
| proseLM-S | ProGen2-small | 1024 | 64 | 16 |
| proseLM-M | ProGen2-medium | 1536 | 96 | 16 |
| proseLM-L | ProGen2-large | 2048 | 128 | 16 |
| proseLM-XL | ProGen2-xlarge | 4096 | 256 | 16 |
| proseLM-Base | ProGen2-base | 1536 | 96 | 16 |
| proseLM-Ab | ProGen2-OAS | 1536 | 96 | 16 |

ProseLM models were trained for 5 epochs on the CATH 4.2 dataset and 15 epochs on the PDB dataset with an effective batch size of 64 using the Adam optimizer (37). The learning rate was set to a fixed value of 2e-4. An additional set of models were trained in the same manner but with 0.1 Ã Gaussian noise added to the protein backbone coordinates. The final models were chosen according to validation set loss.

##### Algorithm 5 proseLM

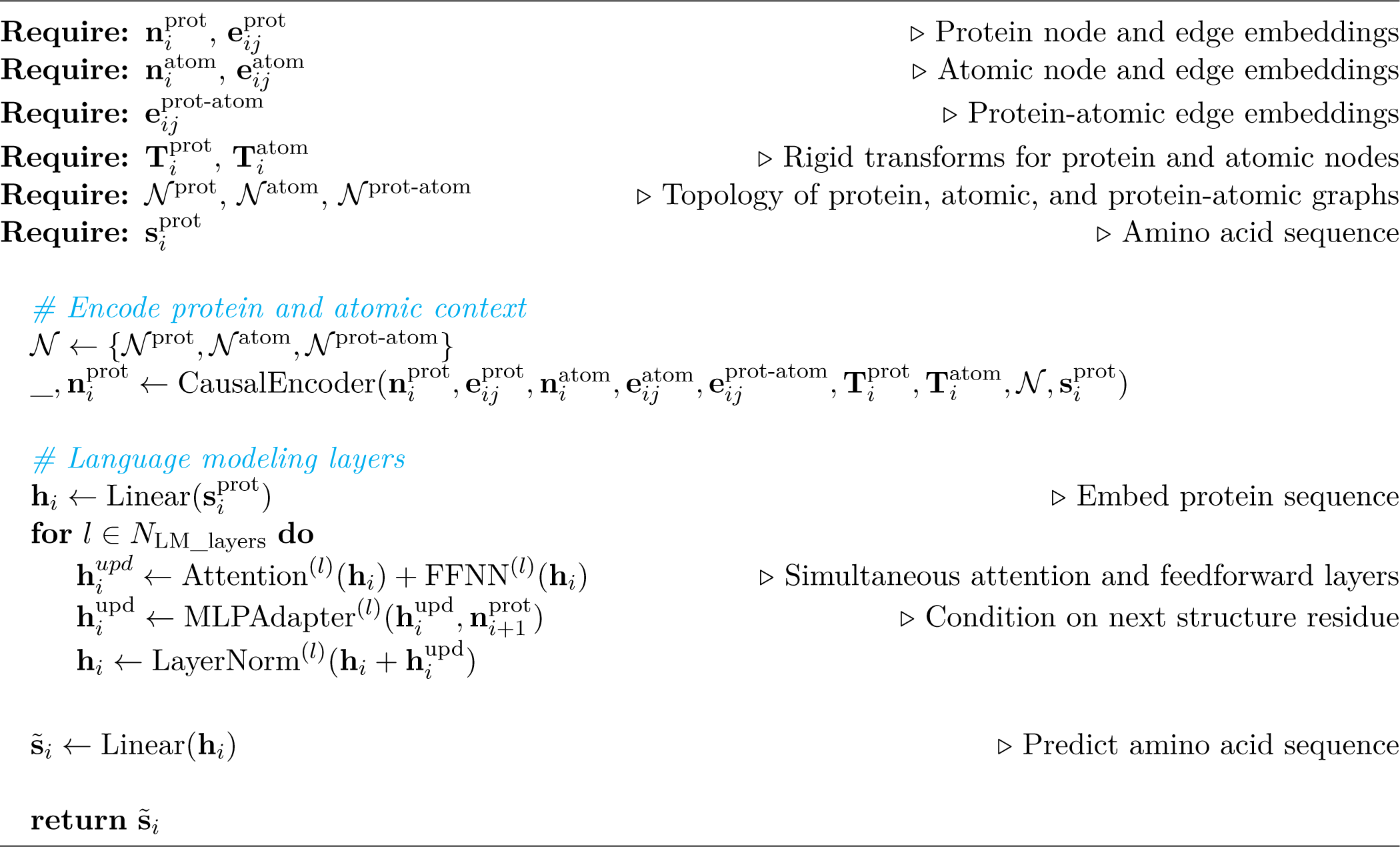

##### Algorithm 6 MLP adapter

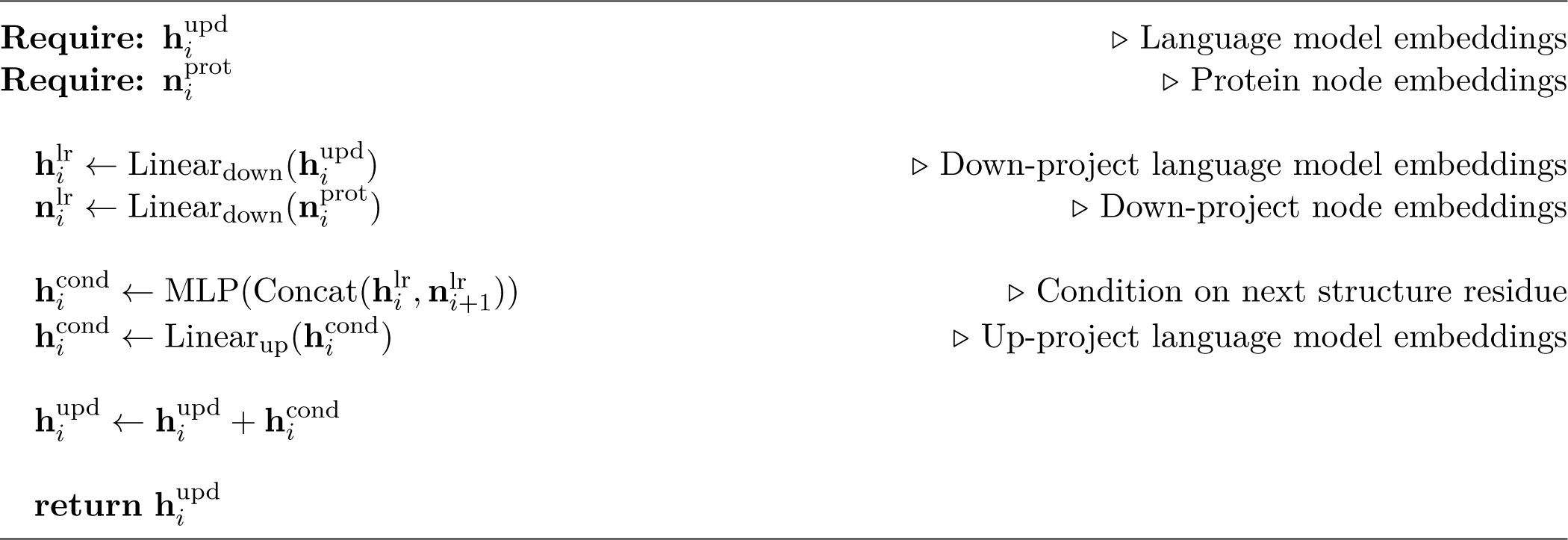

### Comparison to LM-Design

The most conceptually similar method to proseLM is LM-Design, which adapts masked language models for structure-conditioned sequence design (38). While proseLM generates sequences autoregressively for a given backbone structure, LM-Design adopts a strategy more akin to sequence refinement using language models given an initial guess. Architecturally, the conditional adapter of proseLM uses significantly few parameters per layer (80K-300K) than LM-Design (5M per layer). This discrepancy enables proseLM to maintain parameter-efficiency while incorporating adapters after every layer of the language model, whereas LM-Design only incorporates adapters after the final layer. To assess the impact of these architectural differences, we compared proseLM to LM-Design on the CATH 4.2 test set (Table S1). The most apt comparison to the reported LM-Design performance is proseLM-M, which contains a similar number of pre-trained language model parameters. We found that proseLM-M achieves lower perplexity, while LM-Design achieves higher native sequence recovery. This discrepancy likely arises from the different modeling objectives, with proseLM sampling sequences autoregressively and LM-Design iteratively updating sequences by sampling from position-wise marginal distributions. Autoregressive modeling directly captures the co-evolution between pairs of residues, which is critical for protein function, but may yield designs with lower native sequence recovery.

### Design of genome editors

#### Nuclease optimization

To design optimized SpCas9 nucleases with proseLM, we trained proseLM-Base by adapting the ProGen2-base model and training on the PDB dataset. This was necessary in order to model the complete 1,368-residue SpCas9 sequence, which extends beyond the maximum length supported by other ProGen2 and proseLM models. For design, we utilized a multi-state conditioning strategy, selecting two structures for conditioning that represented the binary (PDB ID 4ZT0) and catalytic (PDB ID 7Z4J) states of SpCas9. To account for missing residues in the structures, we used AlphaFold2 to predict the structures with each state provided as a template. The predicted structures were aligned with sub-Å RMSD into the experimental complexes, yielding structurally complete representations of the binary and catalytic states. Using these structures, we designed 1,600 sequences (*T* = 1.0). All sequences were had the PAM-interacting domain and known catalytic residues fixed to their natural identities.

To generate sequences closer to the natural SpCas9 sequence, we introduced positional residue frequencies derived from evolutionary and experimental data. Evolutionary data was obtained from a multiple-sequence alignment of phylogenetically related Cas9 proteins, which were used to create a position-specific scoring matrix (PSSM). Experimental data was obtained from a deep mutational scanning study of SpCas9, which measured the on- and off-target impacts of single amino acid mutations (39). We combined the PSSM and normalized DMS data to create a amino acid bias term for each residue in the SpCas9 sequence. This bias was added to the logits of proseLM-Base and used to generate an additional 4,400 sequences (*T* = 0.1). We selected the seven designs from this set for experimental characterization according to the criteria used for OpenCRISPR (16).

#### Base editor optimization

For base editor optimization, we selected a deaminase domain previously designed using protein language models fine-tuned on the TadA family (16). We predicted the structure of the deaminase dimer using the ABE8e structure (PDB ID 6VPC) as a template, then aligned our predicted structure into the functional state of the ABE8e complex with sub-Å RMSD. We divided our designs across two strategies, focusing separately on active site or non-functional scaffolding residues. For active site designs, we selected all residues within 5 Å of the single-stranded DNA or Cas9 nuclease, excluding positions at the termini or within 5 Å of the dimeric interface, and kept all other residues fixed. For non-active site designs, we selected all residues further than 5 Å from the single-stranded DNA or Cas9 nuclease and further than 5 Å from the dimeric interface, and kept all other residues fixed. When generating designs, we provided the fixed residues to the causal encoder as context by reordering the residue indices, effectively conditioning proseLM on future positions. We generated 300 active site designs (*T* = 1.0) and 200 non-active site designs (*T* = 0.5) with proseLM-S. The 40 best designs from each strategy according to perplexity were selected for experimental characterization.

### Design of therapeutic antibodies

#### Nivolumab optimization

To optimize the binding affinity of the therapeutic antibody nivolumab, we used the crystal structure of nivolumab bound to PD-1 (PDB ID 5WT9). We divided our designs across two strategies, focusing separately on the complementarity-determining regions (CDRs) and the framework regions. For CDR-directed designs, we enumerated all possible single- and double-mutations to residues within 8 Å of the antigen, excluding mutations to or from cysteine or proline. In total, this set included 414,477 variants with mutations across 54 positions. For framework-directed designs, we generated designs with conditioning on the CDRs by fixing residues within 6 Å of the antigen. We generated 2,000 designs each from proseLM-Ab and proseLM-Base (*T* = 1.0). Half of the sequences had the heavy chain designed first, with the light chain successively designed based on the designed heavy, and the other half used the reverse chain order. We selected 55 CDR-directed and 40 framework-directed designs according to an ensemble of proseLM models (causal encoder, proseLM-S, proseLM-Base, proseLM-XL, and proseLM-Ab), using the product of perplexities as a selection criterion. For CDR-directed designs, we selected 15 single mutations and 40 double mutations. For framework-directed designs, we selected 20 designs with 1-10 mutations and 20 designs with 11-20 mutations. For each strategy, we ensured that no particular mutation appeared more than ten times in the final set. In total, 95 designs were selected for experimental characterization.

#### Secukinumab diversification

For diversification of the therapeutic antibody secukinumab, we used the crystal structure of secukinumab bound to IL-17A (PDB ID 6WIO). We generated full heavy and light chain variable fragments with 2,000 designs each from proseLM-Ab and proseLM-Base (*T* = 1.0). Half of the sequences had the heavy chain designed first, with the light chain successively designed based on the designed heavy, and the other half used the reverse chain order. From each model’s designs, we selected 16 designs with 1-24 mutations, 16 designs with 25-29 mutations, and 16 designs with 30-34 mutations according to an ensemble of proseLM models (causal encoder, proseLM-S, proseLM-Base, proseLM-XL, and proseLM-Ab), using the product of perplexities as a selection criterion In total, 96 designs were selected for experimental characterization.

### Characterization of genome editors

#### DNA oligonucleotides and plasmid assembly

Oligonucleotides used in this study were synthesized by IDT with standard desalting. All natural and AI-generated nuclease and deaminase sequences were purchased as synthetic gene fragments (Twist Bioscience), human codon-optimized using the Twist codon optimization tool, and cloned into CMV-driven expression plasmids using HiFi DNA Assembly (New England Biolabs). Single-guide RNA (sgRNA) sequences were cloned into a human U6 (hU6)-driven expression plasmid that also contains a CMV-driven GFP transfection reporter using HiFi DNA Assembly (New England Biolabs). All plasmids were sequence-verified by whole plasmid Nanopore sequencing (Primordium) prior to downstream applications.

#### HEK293T cell culture and transient transfection

HEK293T cells (ATCC) were cultured at 37°C and 5% (v/v) CO2 in high glucose DMEM with 4 mM L-glutamine, 1 mM sodium pyruvate and phenol red pH indicator (Gibco), supplemented with 10% FBS and 1X penicillin-streptomycin. 24 hours prior to transfection, cells were seeded at a density of 1x103 cells/well in 96-well tissue culture-treated plates (Nunc™ Edge™, Thermo Fisher Scientific).

For each transfection well, 50 ng of sgRNA plasmid and 50 ng of nuclease or base editor-expressing plasmid were added to 5 *µ*L of Opti-MEM (Gibco). 0.2 *µ*L of TransIT®-2020 transfection reagent (Mirus Bio) was diluted into 4 *µ*L of Opti-MEM. Plasmid and TransIT®-2020 mixtures were combined, incubated for 15-30 min at room temperature, and added to HEK293T cells in a dropwise manner. Plates were gently rocked to mix and incubated for 72 hours.

#### Targeted-amplicon sequencing and analysis

Cell lysates were generated by washing the cells with 1X PBS and adding 25 *µ*L of lysis buffer (100 mM Tris-HCl, pH 7.5; 0.05% SDS; 25 *µ*g/mL Proteinase K) per well. Plates with lysis buffer were incubated at 37°C for 1 hour, and then 25 *µ*L of nuclease-free water was added per well. The lysates were transferred to 96-well PCR plates and boiled at 98°C for 15 min. Locus-specific primers were used to amplify regions of interest from cell lysates by PCR (Q5® High-Fidelity DNA Polymerase). After PCR, the resulting amplicons were purified (Mag-Bind® RxnPure Plus, Omega Bio-tek), DNA yields were quantified (QuantiFluor® kit, Promega), and DNA concentrations were normalized to 2 ng/100 bp of amplicon length and submitted for Sanger sequencing with the appropriate forward PCR primer. We quantified the activity of base editors and nucleases using the software tools BEAT (40) and Synthego ICE v1.2.0 (41), respectively, with default parameters.

### Characterization of antibodies

#### Antibody production

Antibody heavy and light chain variable fragment sequences were synthesized as gene fragments and cloned into the pTwist CMV vectors. Nivolumab variants were cloned into IgG4 (heavy chain) and IgK (light chain) vectors. Secukinumab variants were cloned into IgG1 (heavy chain) and IgK (light chain) vectors. Antibodies were expressed in 1mL HEK293 mammalian cultures and the supernatant was used for binding affinity measurements. Gene synthesis and expression was performed by Twist Bioscience.

#### Binding affinity measurements

Binding studies were performed in HBSTE running buffer (10mM HEPES, 150mM NaCl, 3mM EDTA, 0.0% Tween-20) at 25degC. Six-point antigen dilution series were prepared in running buffer starting at 200 nM with a 2.5-fold serial dilution (200-2.0nM). Antibodies were first captured on a Carterra HC30M sensor chip with immobilized goat anti-human Fc pAb. Following 8-10 buffer injection cycles, increasing antigen concentrations were injected over the Ab-captured surfaces with a 5 min of association phase and a 10 min of dissociation phase. The chip surface was finally regenerated and antibodies were recaptured for subsequent antigen binding studies. Double reference subtracted data containing antigen binding at varying concentrations were globally fit using 1:1 binding model. Binding affinity measurements and analyses were performed by Twist Bioscience.

## Supplementary information

**Table S1.** CATH 4.2 benchmark. Comparison of recent protein design models on the CATH 4.2 test set from (2). Perplexity is reported over all residues in the test set. † indicates results cited from (22). ‡ indicates results cited from (38).

| Model | Train / Total Param. | Perplexity (↓) | Median Recovery (% , ↑) |
| --- | --- | --- | --- |
| †Structured Transformer (2) | 1.6M / 1.6M | 6.63 | 35.82 |
| †GVP-GNN (42) | 1.0M / 1.0M | 5.36 | 39.47 |
| †ProteinMPNN (3) | 1.9M / 1.9M | 4.61 | 45.96 |
| †PiFold (22) | 6.6M / 6.6M | 4.55 | 51.66 |
| ‡ProteinMPNN-CMLM (38) | 1.9M / 1.9M | 5.03 | 48.62 |
| ‡LM-Design (ProteinMPNN-CMLM) (38) | 6.9M / 659M (1.0%) | 4.63 | 53.26 |
| ‡LM-Design (PiFold) (38) | 11.9M / 664M (1.7%) | 4.52 | 55.65 |
| Causal encoder | 4.1M / 4.1M | 4.76 | 47.24 |
| proseLM-S | 5.1M / 156M (3.3%) | 3.89 | 49.00 |
| proseLM-Base | 7.3M / 772M (0.9%) | 3.34 | 49.6 |
| proseLM-M | 7.3M / 772M (0.9%) | 3.29 | 50.00 |
| proseLM-L | 10.8M / 2.8B (0.4%) | 3.23 | 50.34 |
| proseLM-XL | 13.3M / 6.5B (0.2%) | 2.67 | 50.83 |

**Table S2.** PDB benchmark. Comparison of proseLM models on PDB test set derived from (3). Metrics are for proteins that are part of a complex or have interactions with some non-protein entity. Performance is reported for both backbone-only and full-context inputs (backbone / full). For protein complexes, all other chains are provided as context for the target chain.

| Model | Train / Total Param. | Mean Perplexity (↓) | Median Recovery (% , ↑) |
| --- | --- | --- | --- |
| Causal encoder | 4.2M / 4.2M | 4.17 / 3.64 | 54.36 / 58.15 |
| proseLM-S | 988K / 155M (0.6%) | 3.77 / 3.34 | 55.63 / 59.57 |
| proseLM-M | 3.2M / 772M (0.4%) | 3.32 / 3.00 | 57.00 / 61.20 |
| proseLM-L | 6.6M / 2.8B (0.2%) | 3.39 / 3.05 | 56.89 / 61.53 |
| proseLM-XL | 9.2M / 6.5B (0.1%) | 2.80 / 2.57 | 59.58 / 64.41 |

**Table S3.** Sequence recovery near non-protein context. Comparison of proseLM models on PDB test set derived from (3). Metrics are for residues within 5 Å of nucleic acids, ligands, or ions. Performance is reported for both backbone-only and full-context inputs (backbone / full).

| Model | Median Recovery (% , ↑) |  |  |
| --- | --- | --- | --- |
|  | Nucleic Acid | Ligand | Ion |
| Causal encoder | 44.44 / 54.90 | 50.00 / 64.00 | 46.15 / 66.67 |
| proseLM-S | 48.94 / 58.06 | 57.14 / 70.59 | 60.00 / 76.00 |
| proseLM-M | 64.24 / 70.87 | 66.67 / 75.93 | 70.97 / 82.61 |
| proseLM-L | 62.88 / 71.13 | 66.67 / 76.47 | 70.59 / 82.61 |
| proseLM-XL | 70.59 / 76.10 | 74.19 / 82.35 | 77.78 / 87.50 |

**Table S4.** SAbDab antibody benchmark. Comparison of proseLM models on antibody test set from SAbDab. Sequence recovery is reported as the median of cluster-averaged values for all antigen-bound heavy (n = 361) and light (n = 300) chains in the test set. Performance is reported for both antibody-only and full-complex inputs (antibody / complex).

| Model | Median Recovery (% , ↑) |  |
| --- | --- | --- |
|  | Heavy Fv | Light Fv |
| Causal encoder | 75.90 / 77.37 | 75.68 / 77.48 |
| proseLM-S | 78.56 / 78.94 | 77.57 / 78.06 |
| proseLM-M | 79.91 / 79.96 | 79.61 / 79.44 |
| proseLM-L | 80.71 / 82.04 | 82.14 / 83.17 |
| proseLM-XL | 80.60 / 81.57 | 80.09 / 81.02 |
| proseLM-Ab | 79.80 / 82.42 | 83.11 / 83.02 |

**Table S5.** SAbDab bound antibody benchmark. Comparison of proseLM models on antibody test set from SAbDab. Sequence recovery is reported as the median of cluster-averaged values for all antigen-bound heavy (n = 361) and light (n = 300) chains in the test set. Performance is reported for both antibody-only and full-complex inputs (antibody / complex). CDR loops are defined according to the IMGT numbering scheme.

| Model | Median Recovery (% , ↑) |  |  |  |  |  |  |  |
| --- | --- | --- | --- | --- | --- | --- | --- | --- |
|  | Heavy Fr | CDR H1 | CDR H2 | CDR H3 | Light Fr | CDR L1 | CDR L2 | CDR L3 |
| Causal encoder | 83.23 / 82.73 | 67.22 / 69.23 | 55.56 / 60.00 | 50.00 / 60.00 | 81.01 / 82.28 | 65.00 / 66.67 | 62.50 / 62.50 | 56.25 / 59.09 |
| proseLM-S | 85.42 / 84.70 | 69.23 / 72.48 | 60.00 / 63.65 | 52.51 / 57.70 | 82.28 / 82.28 | 63.64 / 66.20 | 62.50 / 68.75 | 56.36 / 60.42 |
| proseLM-M | 87.50 / 87.57 | 69.23 / 69.23 | 60.00 / 62.60 | 52.66 / 58.82 | 84.81 / 84.21 | 66.67 / 66.67 | 66.67 / 75.00 | 57.78 / 60.00 |
| proseLM-L | 88.62 / 88.10 | 69.41 / 74.03 | 60.00 / 66.67 | 52.63 / 62.83 | 86.00 / 86.08 | 68.18 / 68.18 | 75.00 / 75.00 | 65.08 / 66.67 |
| proseLM-XL | 88.39 / 88.11 | 69.23 / 70.19 | 60.00 / 68.93 | 50.00 / 60.56 | 84.54 / 85.53 | 69.23 / 71.10 | 71.43 / 75.00 | 59.26 / 62.50 |
| proseLM-Ab | 89.09 / 89.70 | 69.23 / 71.43 | 59.44 / 63.33 | 50.00 / 60.41 | 86.71 / 86.71 | 66.67 / 72.73 | 75.00 / 75.00 | 62.50 / 66.67 |

**Fig. S1.**
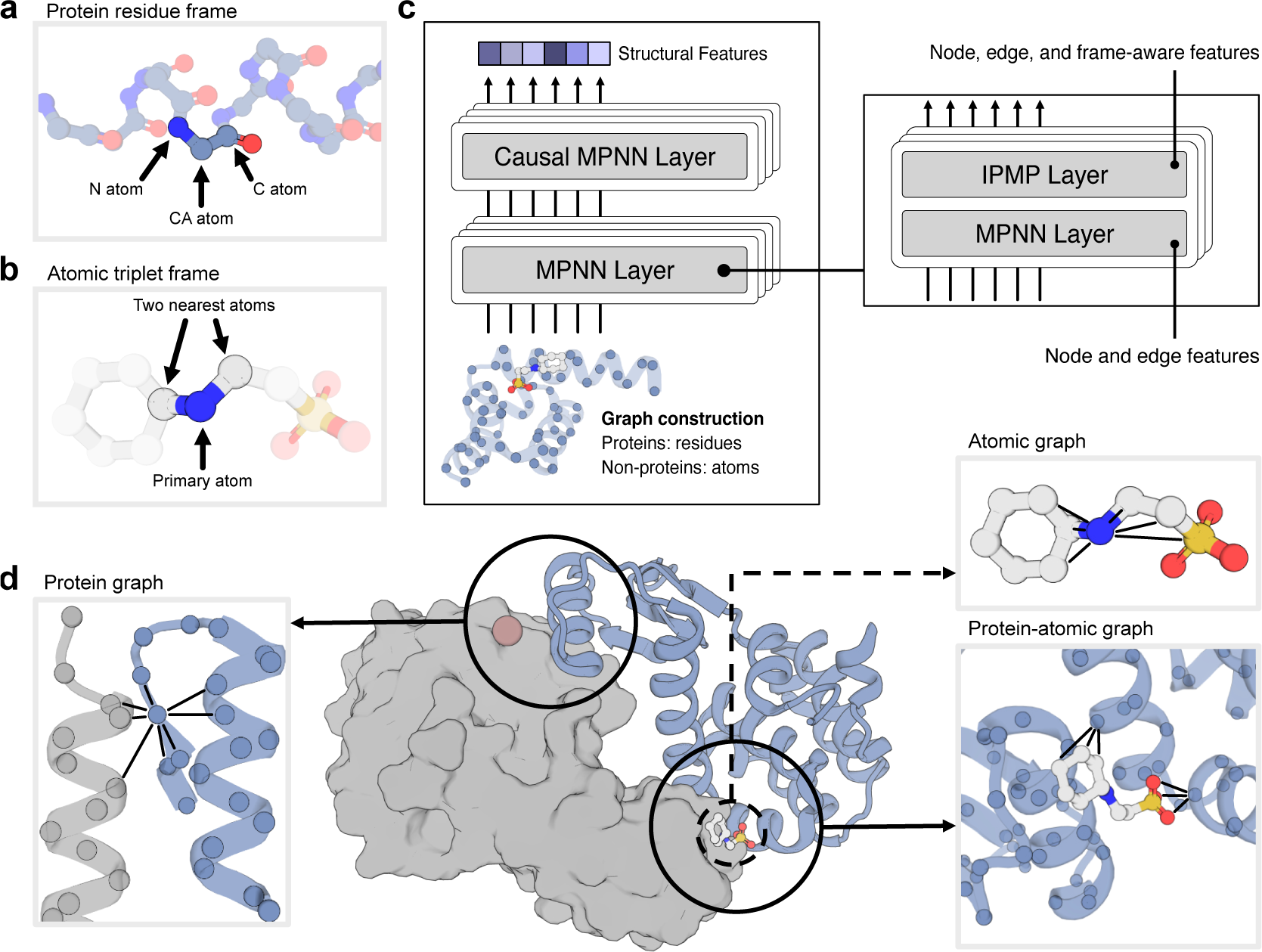
Visualization of protein and atomic graph components. (a) Each protein residue is represented as a rigid body frame with the C*_α_* atom at the origin and the frame’s orientation defined by the positions of the N and C atoms. (b) Non-protein atoms are represented as rigid body frames with the primary atom at the origin and the two closest atoms defining the frame’s orientation. (c) Protein and atomic graphs are processed by alternating MPNN and IPMP (20) layers. MPNN layers operate on the graph nodes and edges, while IPMP layers additionally incorporate frame-based geometric features. (d) Visualization of the edge connectivity for the protein-only residue graph (left), atomic-only graph (upper right), and protein-atomic graph (lower right). Black lines illustrate the connectivity between nodes in the graph.

**Fig. S2.**
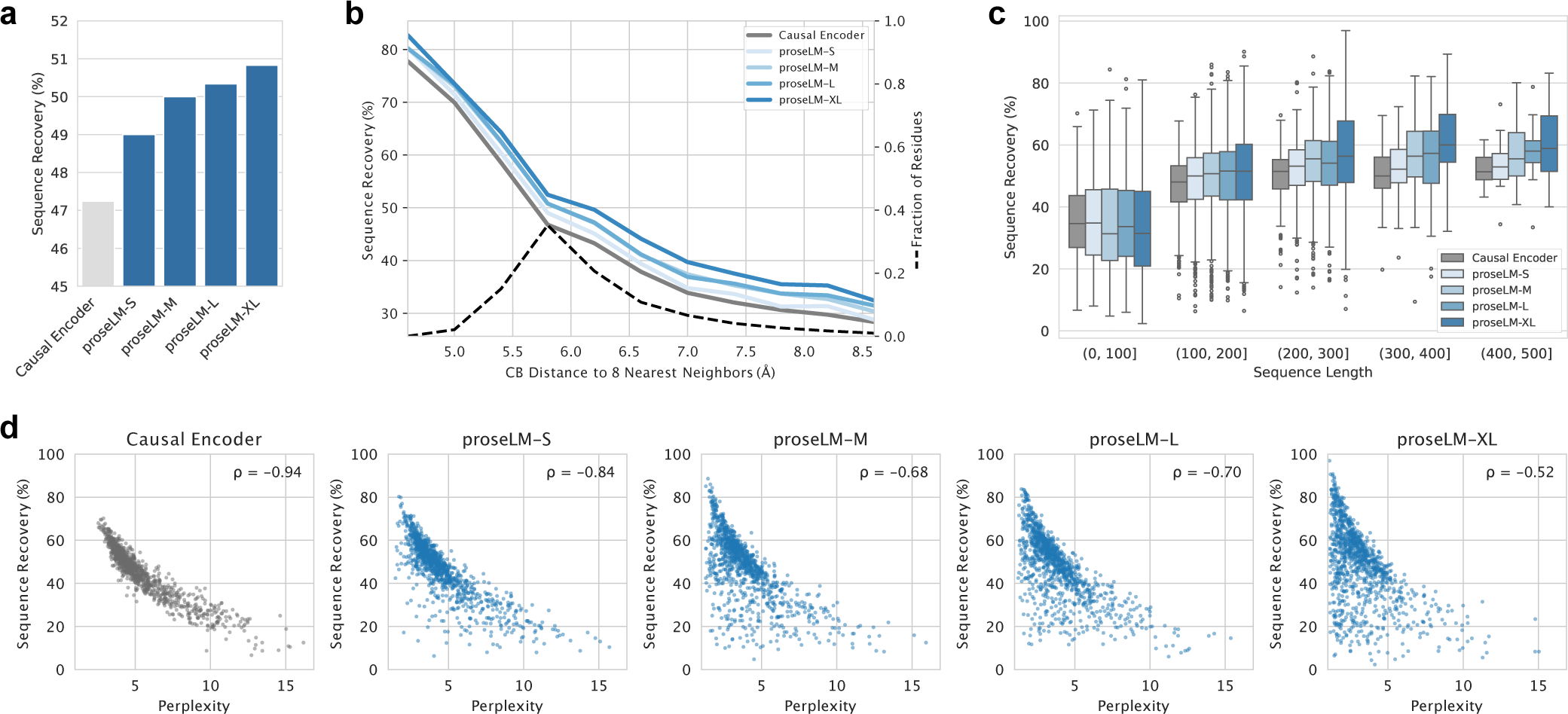
Performance of proseLM models on CATH 4.2 benchmark. Evaluation of proseLM model performance on the CATH 4.2 test set curated by Ingraham et al. (2). Aggregate sequence recovery values are reported as the median across all proteins (or subsets when appropriate). (a) Recovery of native sequence residues for proseLM models. (b) Recovery of native sequence residues for proseLM models binned by residue burial, calculated as the average C*_β_* distance to the nearest eight neighbors (lower is more buried). All models achieve highest rates of native sequence recovery among buried residues and reduced recovery at less-buried surface positions. (c) Recovery of native sequence residues for proseLM models, binned by sequence length. Longer sequences exhibit higher rates of native sequence recovery, with larger proseLM models having particularly high recovery for large proteins. (d) Relationship between perplexity and sequence recovery for proseLM models. Spearman correlation coefficients are reported for each model. Larger models show less correlation between perplexity and sequence recovery.

**Fig. S3.**
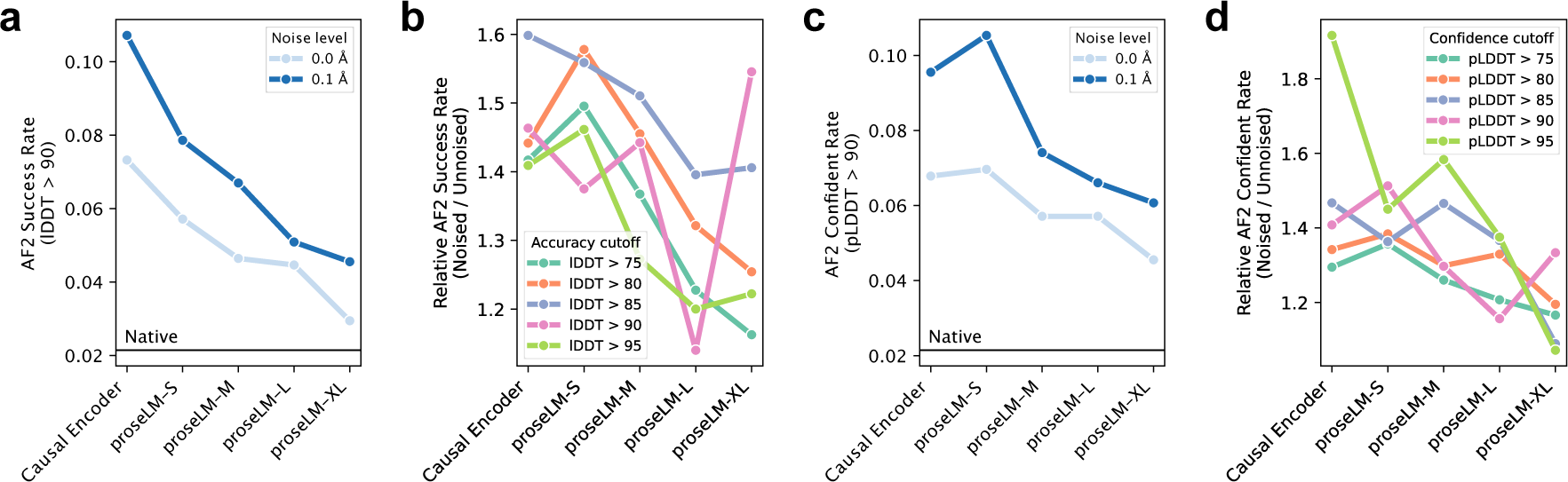
Impact of coordinate noise on single-sequence structure prediction. Evaluation of proseLM model performance on the CATH 4.2 test set with and without Gaussian noise added to coordinates. Structure prediction accuracy and confidence are for single-sequence predictions using AlphaFold2 (9). (a) Fraction of designed sequences that are predicted to successfully recapitulate the input structure (lDDT > 90). Models trained with 0.1 Å of Gaussian noise added to coordinates achieve higher structure prediction success. Larger models approach the lower level of structure prediction success of the native sequences (Horizontal line) (b) Structure prediction success rates for noised models relative to un-noised for several lDDT thresholds. Larger models show less sensitivity to coordinate noise (lower relative success rates). (c) Fraction of designed sequences that yield highly confident structure predictions (pLDDT > 90). Models trained with coordinate noise yield high-confidence structures more frequently. Larger models approach the lower level of confident structure prediction rates for the native sequences (Horizontal line). (d) Confidence in structure prediction for noised models relative to un-noised for several pLDDT thresholds. Larger models show less sensitivity to coordinate noise (lower relative rates of confident predictions).

**Fig. S4.**
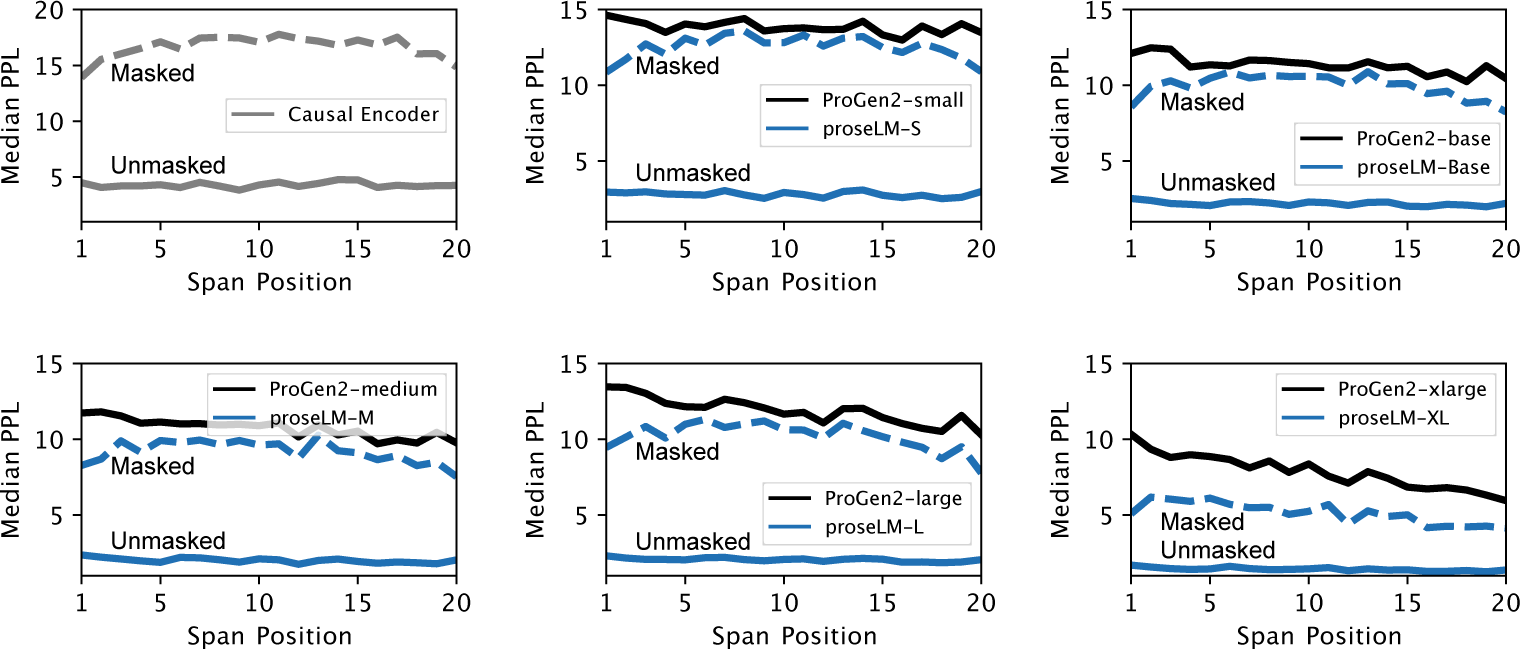
Perplexity of masked structural spans. Evaluation of proseLM model perplexity on contiguous structurally masked spans of twenty residues. Perplexity for structurally masked residues (dashed lines) increases significantly over the same residues without masking (solid lines). Compared to the respective ProGen2 models, proseLM models achieve lower perplexity for structurally masked residues, indicating that the surrounding context is effectively incorporated when predicting for masked positions.

**Fig. S5.**
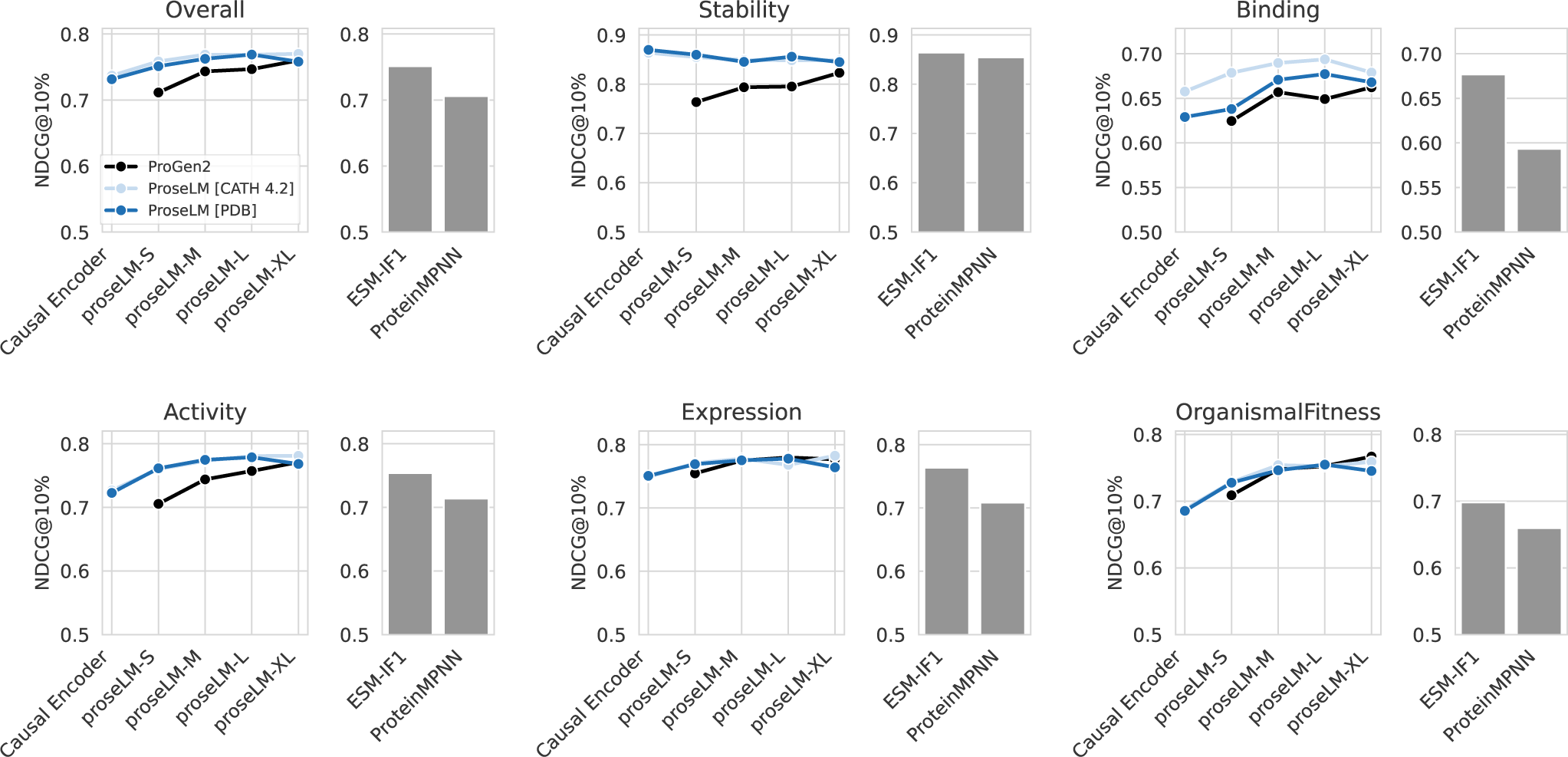
Fitness prediction measured by normalized discounted cumulative gain. Comparison of proseLM and recent structure-conditioned sequence design models (ESM-IF1 and ProteinMPNN) for prediction of mutational fitness landscapes. Performance is reported as normalized discounted cumulative gain for the top ten percent of samples according to experimental fitness (NDCG10%). The metric is relevant for protein design tasks, where it is important to accurately prioritize the highest fitness sequences for experimental characterization.

**Fig. S6.**
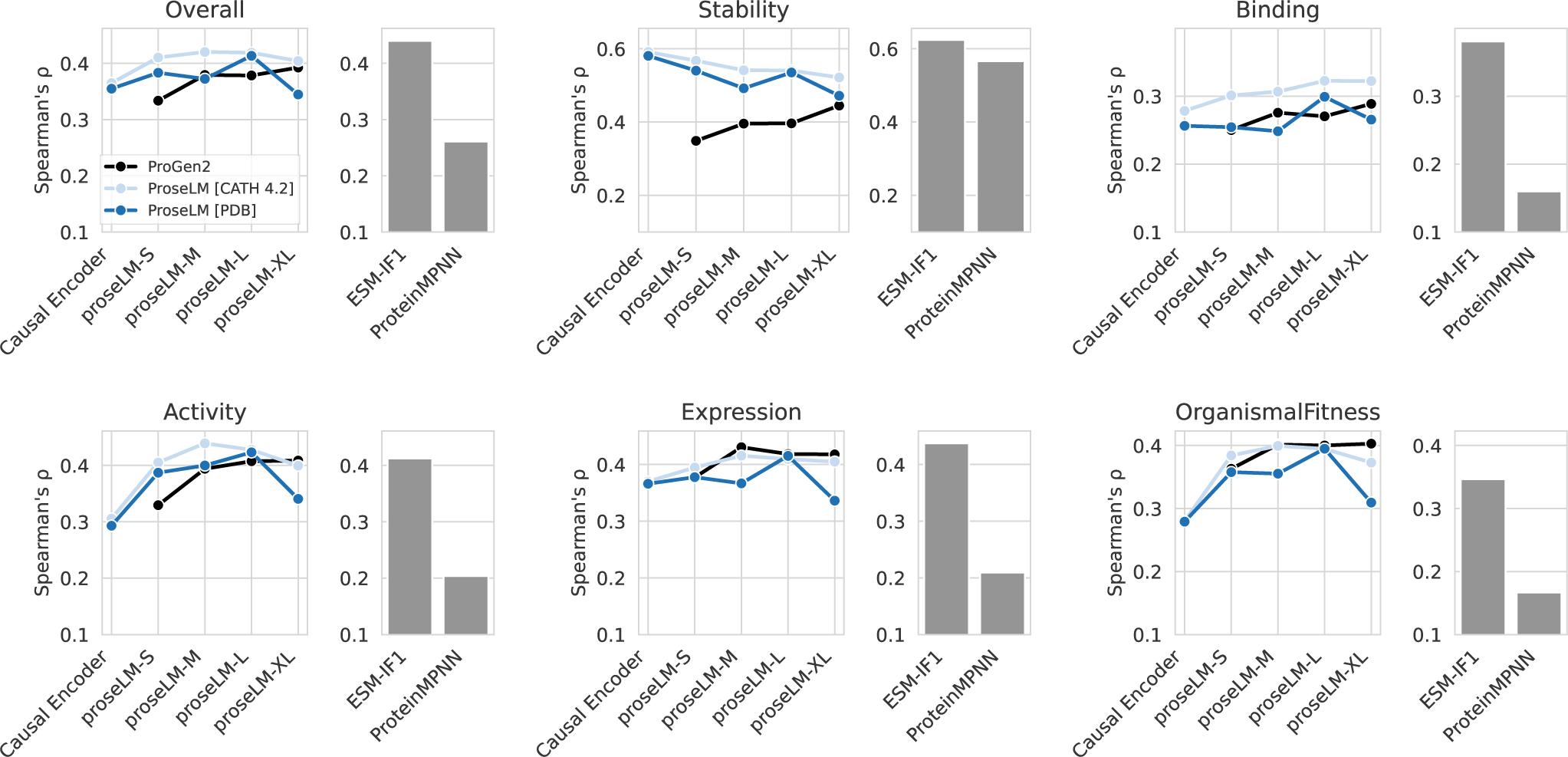
Fitness prediction measured by Spearman’s rank correlation coefficient. Comparison of proseLM and recent structure-conditioned sequence design models (ESM-IF1 and ProteinMPNN) for prediction of mutational fitness landscapes. Performance is reported as Spearman’s rank correlation coefficients. The metric indicates how well model scores rank sequences according to fitness across the entire dataset.

**Fig. S7.**
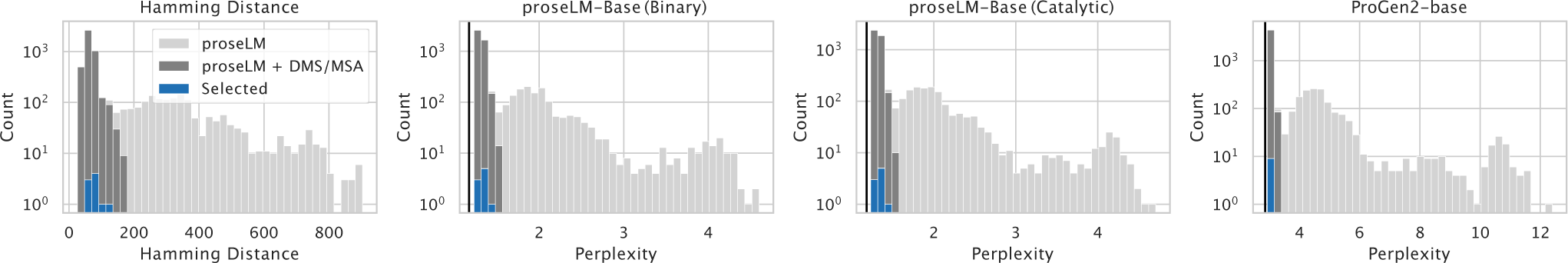
Model scores for SpCas9 variants. Mutational hamming distance from SpCas9 and proseLM model perplexities for designed SpCas9 variants. Mutation distances and model scores are shown for generations directly from proseLM (light gray), generations from proseLM augmented with position-specific residue propensities from deep mutational scans and multiple-sequence alignments (dark gray), and the subset selected for experimental validation (blue). Model scores for SpCas9 are indicated by vertical black lines.

**Fig. S8.**
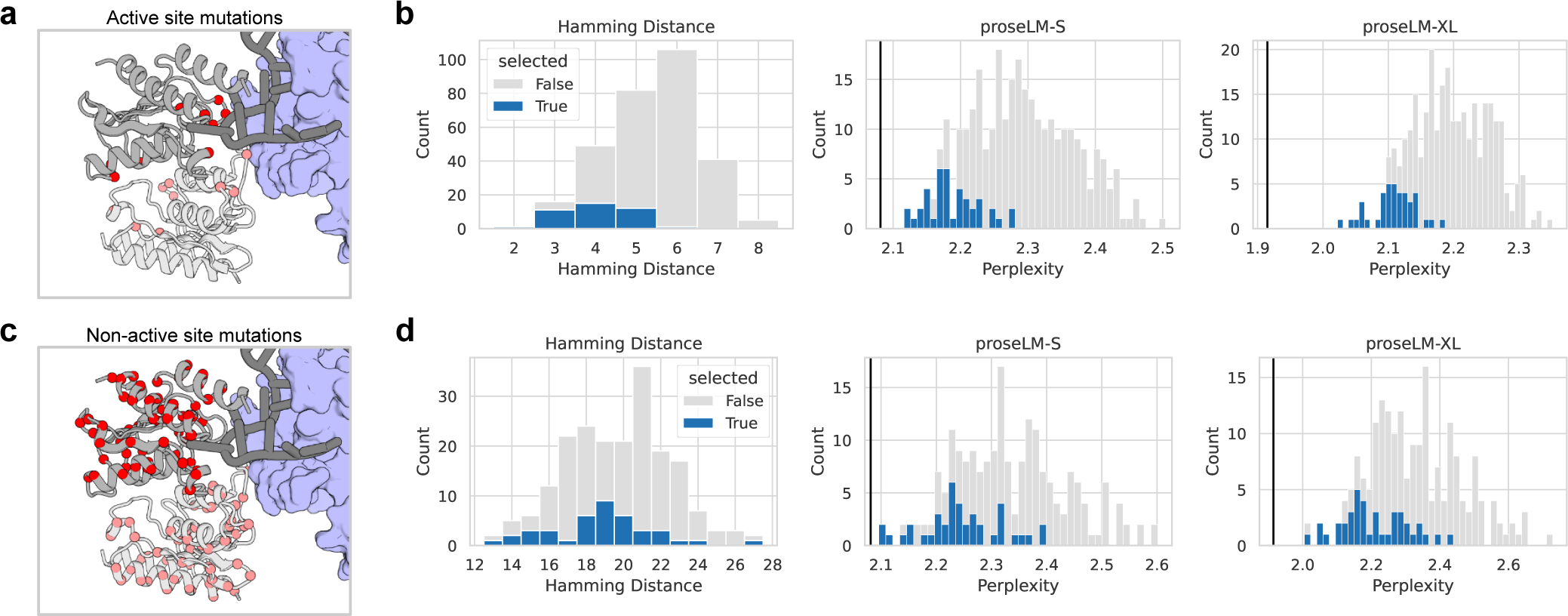
Model scores for adenine base editor variants. Mutational hamming distance from parental deaminase and proseLM model perplexities for designed deaminase variants. Mutation distances and model scores for the complete set of generations and the subset selected for experimental validation are shown in gray and blue, respectively. Model scores for the parental deaminase are indicated by vertical black lines. (a) Positions of all mutated residues among selected active site variants (red spheres). (b) Mutation and score distributions for active site variants. (c) Positions of all mutated residues among selected non-active site variants (red spheres). (d) Mutation and score distributions for non-active site variants.

**Fig. S9.**
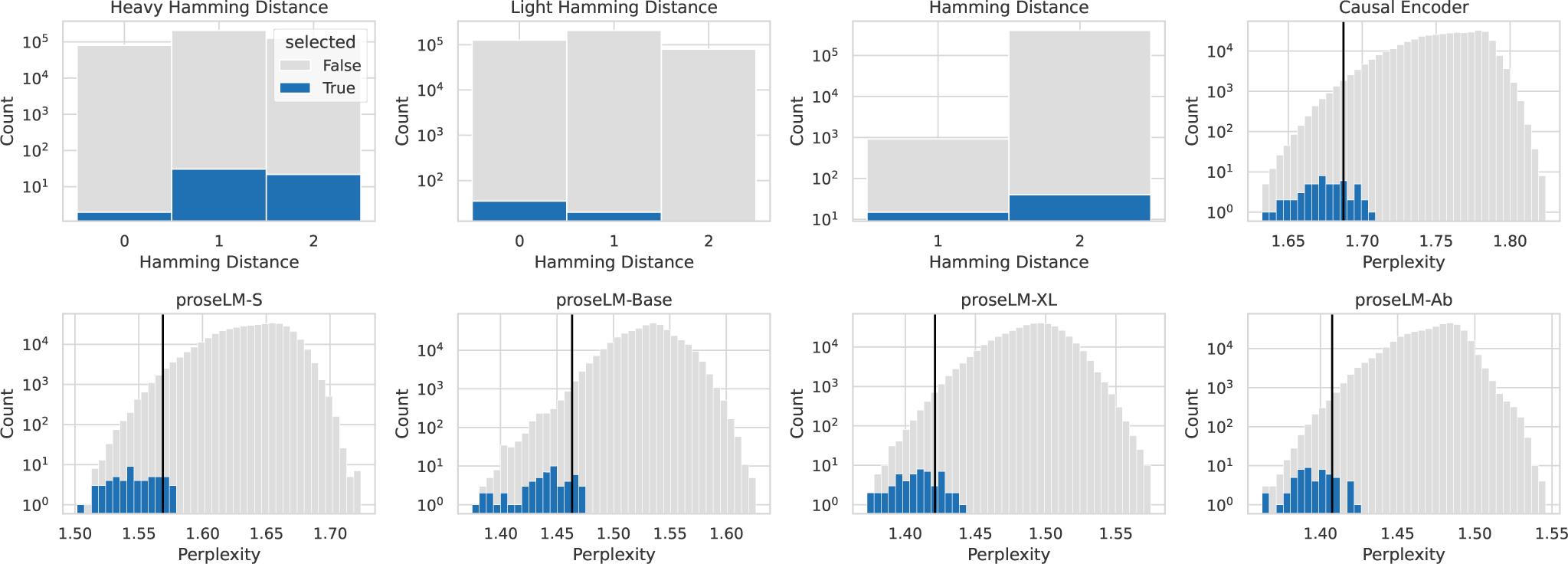
Model scores for nivolumab CDR variants. Mutational hamming distance from parental nivolumab and proseLM model perplexities for designed nivolumab CDR loop variants. Mutation distances and model scores for the complete set of generations and the subset selected for experimental validation are shown in gray and blue, respectively. Model scores for nivolumab are indicated by vertical black lines. Y-axis is shown on a log scale.

**Fig. S10.**
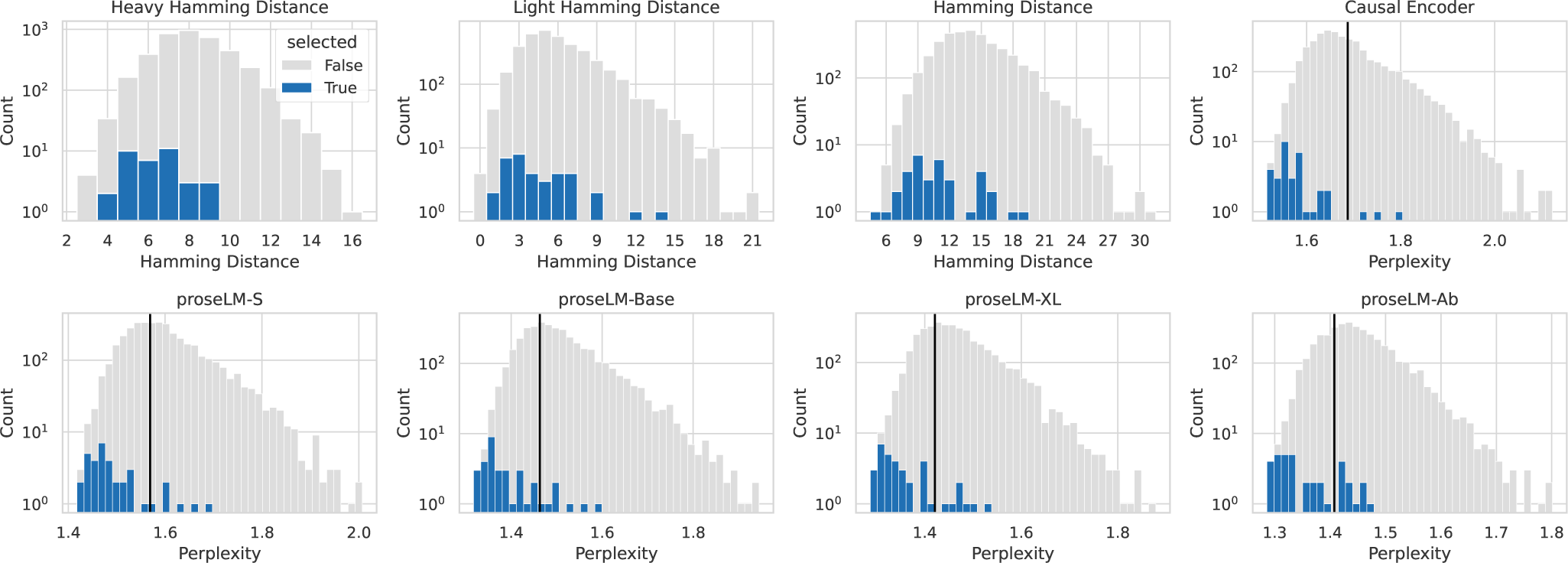
Model scores for nivolumab framework variants. Mutational hamming distance from parental nivolumab and proseLM model perplexities for designed nivolumab framework variants. Mutation distances and model scores for the complete set of generations and the subset selected for experimental validation are shown in gray and blue, respectively. Model scores for nivolumab are indicated by vertical black lines. Y-axis is shown on a log scale.

**Fig. S11.**
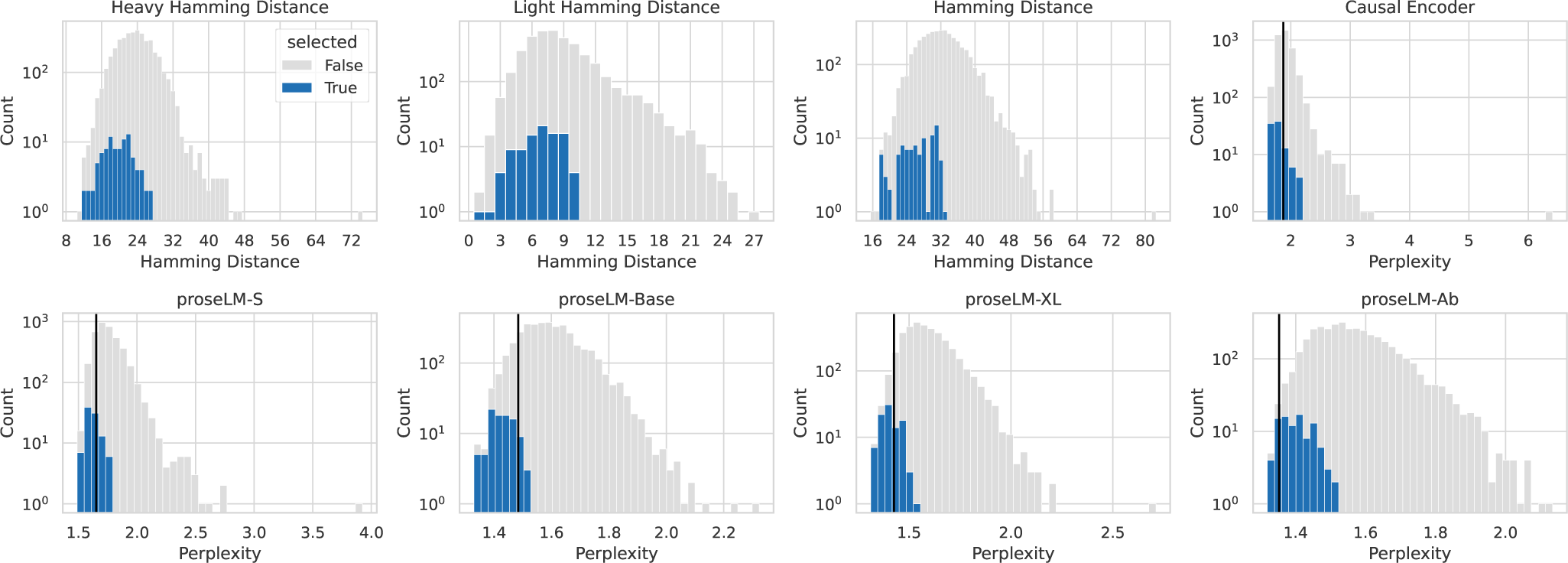
Model scores for diversified secukinumab variants. Mutational hamming distance from parental secukinumab and proseLM model perplexities for designed secukinumab variants. Mutation distances and model scores for the complete set of generations and the subset selected for experimental validation are shown in gray and blue, respectively. Model scores for secukinumab are indicated by vertical black lines. Y-axis is shown on a log scale.

## References

1. Andrew Leaver-Fay, Michael Tyka, Steven M Lewis, Oliver F Lange, James Thompson, Ron Jacak, Kristian W Kaufman, P Douglas Renfrew, Colin A Smith, Will Sheffler, et al. Rosetta3: an object-oriented software suite for the simulation and design of macromolecules. In *Methods in enzymology*, volume 487, pages 545–574. Elsevier, 2011.

2. John Ingraham, Vikas Garg, Regina Barzilay, and Tommi Jaakkola. Generative models for graph-based protein design. Advances in neural information processing systems, 32, 2019.

3. Justas Dauparas, Ivan Anishchenko, Nathaniel Bennett, Hua Bai, Robert J Ragotte, Lukas F Milles, Basile IM Wicky, Alexis Courbet, Rob J de Haas, Neville Bethel, et al. Robust deep learning–based protein sequence design using proteinmpnn. Science, 378(6615):49–56, 2022.

4. Chloe Hsu, Robert Verkuil, Jason Liu, Zeming Lin, Brian Hie, Tom Sercu, Adam Lerer, and Alexander Rives. Learning inverse folding from millions of predicted structures. In International Conference on Machine Learning, pages 8946–8970. PMLR, 2022.

5. BIM Wicky, LF Milles, A Courbet, RJ Ragotte, J Dauparas, E Kinfu, S Tipps, RD Kibler, M Baek, F DiMaio, et al. Hallucinating symmetric protein assemblies. Science, 378(6615):56–61, 2022.

6. Kiera H Sumida, Reyes Núñez-Franco, Indrek Kalvet, Samuel J Pellock, Basile IM Wicky, Lukas F Milles, Justas Dauparas, Jue Wang, Yakov Kipnis, Noel Jameson, et al. Improving protein expression, stability, and function with proteinmpnn. Journal of the American Chemical Society, 146(3):2054–2061, 2024.

7. Florian Praetorius, Philip JY Leung, Maxx H Tessmer, Adam Broerman, Cullen Demakis, Acacia F Dishman, Arvind Pillai, Abbas Idris, David Juergens, Justas Dauparas, et al. Design of stimulus-responsive two-state hinge proteins. bioRxiv, pages 2023–01, 2023.

8. Helen M Berman, John Westbrook, Zukang Feng, Gary Gilliland, Talapady N Bhat, Helge Weissig, Ilya N Shindyalov, and Philip E Bourne. The protein data bank. Nucleic acids research, 28(1): 235–242, 2000.

9. John Jumper, Richard Evans, Alexander Pritzel, Tim Green, Michael Figurnov, Olaf Ronneberger, Kathryn Tunyasuvunakool, Russ Bates, Augustin Žídek, Anna Potapenko, et al. Highly accurate protein structure prediction with alphafold. Nature, 596(7873):583–589, 2021.

10. Roshan Rao, Joshua Meier, Tom Sercu, Sergey Ovchinnikov, and Alexander Rives. Transformer protein language models are unsupervised structure learners. In International Conference on Learning Representations, 2020.

11. Zeming Lin, Halil Akin, Roshan Rao, Brian Hie, Zhongkai Zhu, Wenting Lu, Nikita Smetanin, Robert Verkuil, Ori Kabeli, Yaniv Shmueli, et al. Evolutionary-scale prediction of atomic-level protein structure with a language model. Science, 379(6637):1123–1130, 2023.

12. Daniel Hesslow, Niccoló Zanichelli, Pascal Notin, Iacopo Poli, and Debora Marks. Rita: a study on scaling up generative protein sequence models. arXiv preprint arXiv:2205.05789, 2022.

13. Erik Nijkamp, Jeffrey A Ruffolo, Eli N Weinstein, Nikhil Naik, and Ali Madani. Progen2: exploring the boundaries of protein language models. Cell Systems, 2023.

14. Ali Madani, Ben Krause, Eric R Greene, Subu Subramanian, Benjamin P Mohr, James M Holton, Jose Luis Olmos Jr, Caiming Xiong, Zachary Z Sun, Richard Socher, et al. Large language models generate functional protein sequences across diverse families. Nature Biotechnology, pages 1–8, 2023.

15. Geraldene Munsamy, Ramiro Illanes-Vicioso, Silvia Funcillo, Ioanna T Nakou, Sebastian Lindner, Gavin Ayres, Lesley S Sheehan, Steven Moss, Ulrich Eckhard, Philipp Lorenz, et al. Conditional language models enable the efficient design of proficient enzymes. bioRxiv, pages 2024–05, 2024.

16. Jeffrey A Ruffolo, Stephen Nayfach, Joseph Gallagher, Aadyot Bhatnagar, Joel Beazer, Riffat Hussain, Jordan Russ, Jennifer Yip, Emily Hill, Martin Pacesa, et al. Design of highly functional genome editors by modeling the universe of crispr-cas sequences. bioRxiv, pages 2024–04, 2024.

17. Jonas Pfeiffer, Aishwarya Kamath, Andreas Rücklé, Kyunghyun Cho, and Iryna Gurevych. Adapterfusion: Non-destructive task composition for transfer learning. arXiv preprint arXiv:2005.00247, 2020.

18. Neil Houlsby, Andrei Giurgiu, Stanislaw Jastrzebski, Bruna Morrone, Quentin De Laroussilhe, Andrea Gesmundo, Mona Attariyan, and Sylvain Gelly. Parameter-efficient transfer learning for nlp. In International Conference on Machine Learning, pages 2790–2799. PMLR, 2019.

19. Edward J Hu, Yelong Shen, Phillip Wallis, Zeyuan Allen-Zhu, Yuanzhi Li, Shean Wang, Lu Wang, and Weizhu Chen. Lora: Low-rank adaptation of large language models. arXiv preprint arXiv:2106.09685, 2021.

20. Nicholas Z Randolph and Brian Kuhlman. Invariant point message passing for protein side chain packing. Proteins: Structure, Function, and Bioinformatics, 2024.

21. Christine A Orengo, Alex D Michie, Susan Jones, David T Jones, Mark B Swindells, and Janet M Thornton. Cath–a hierarchic classification of protein domain structures. Structure, 5(8):1093–1109, 1997.

22. Zhangyang Gao, Cheng Tan, and Stan Z Li. Pifold: Toward effective and efficient protein inverse folding. In The Eleventh International Conference on Learning Representations, 2022.

23. Justas Dauparas, Gyu Rie Lee, Robert Pecoraro, Linna An, Ivan Anishchenko, Cameron Glasscock, and David Baker. Atomic context-conditioned protein sequence design using ligandmpnn. Biorxiv, pages 2023–12, 2023.

24. Lucien Krapp, Fernado Meireles, Luciano Abriata, and Matteo Dal Peraro. Context-aware geometric deep learning for protein sequence design. bioRxiv, pages 2023–06, 2023.

25. Pascal Notin, Aaron W Kollasch, Daniel Ritter, Lood van Niekerk, Steffanie Paul, Hansen Spinner, Nathan Rollins, Ada Shaw, Ruben Weitzman, Jonathan Frazer, et al. Proteingym: Large-scale benchmarks for protein design and fitness prediction. bioRxiv, pages 2023–12, 2023.

26. Nicole M Gaudelli, Alexis C Komor, Holly A Rees, Michael S Packer, Ahmed H Badran, David I Bryson, and David R Liu. Programmable base editing of a• t to g• c in genomic dna without dna cleavage. Nature, 551(7681):464–471, 2017.

27. Nicole M Gaudelli, Dieter K Lam, Holly A Rees, Noris M Solá-Esteves, Luis A Barrera, David A Born, Aaron Edwards, Jason M Gehrke, Seung-Joo Lee, Alexander J Liquori, et al. Directed evolution of adenine base editors with increased activity and therapeutic application. Nature biotechnology, 38(7):892–900, 2020.

28. Audrone Lapinaite, Gavin J Knott, Cody M Palumbo, Enrique Lin-Shiao, Michelle F Richter, Kevin T Zhao, Peter A Beal, David R Liu, and Jennifer A Doudna. Dna capture by a crispr-cas9–guided adenine base editor. Science, 369(6503):566–571, 2020.

29. Brian L Hie, Varun R Shanker, Duo Xu, Theodora UJ Bruun, Payton A Weidenbacher, Shaogeng Tang, Wesley Wu, John E Pak, and Peter S Kim. Efficient evolution of human antibodies from general protein language models. Nature Biotechnology, 2023.

30. David Prihoda, Jad Maamary, Andrew Waight, Veronica Juan, Laurence Fayadat-Dilman, Daniel Svozil, and Danny A Bitton. Biophi: a platform for antibody design, humanization, and humanness evaluation based on natural antibody repertoires and deep learning. In *MAbs*, volume 14, page 2020203. Taylor & Francis, 2022.

31. James Dunbar, Konrad Krawczyk, Jinwoo Leem, Terry Baker, Angelika Fuchs, Guy Georges, Jiye Shi, and Charlotte M Deane. Sabdab: the structural antibody database. Nucleic acids research, 42 (D1):D1140–D1146, 2014.

32. Richard W Shuai, Jeffrey A Ruffolo, and Jeffrey J Gray. Iglm: Infilling language modeling for antibody sequence design. Cell Systems, 14(11):979–989, 2023.

33. Mark Hutchinson, Jeffrey A Ruffolo, Nantaporn Haskins, Michael Iannotti, Giuliana Vozza, Tony Pham, Nurjahan Mehzabeen, Harini Shandilya, Keith Rickert, Rebecca Croasdale-Wood, et al. Toward enhancement of antibody thermostability and affinity by computational design in the absence of antigen. mAbs, 16(1):2362775, 2024.

34. Yeqing Lin, Minji Lee, Zhao Zhang, and Mohammed AlQuraishi. Out of many, one: Designing and scaffolding proteins at the scale of the structural universe with genie 2. arXiv preprint arXiv:2405.15489, 2024.

35. Deniz Akpinaroglu, Kosuke Seki, Amy Guo, Eleanor Zhu, Mark JS Kelly, and Tanja Kortemme. Structure-conditioned masked language models for protein sequence design generalize beyond the native sequence space. bioRxiv, pages 2023–12, 2023.

36. Martin Steinegger and Johannes Söding. Clustering huge protein sequence sets in linear time. Nature communications, 9(1):1–8, 2018.

37. Diederik P Kingma and Jimmy Ba. Adam: A method for stochastic optimization. arXiv preprint arXiv:1412.6980, 2014.

38. Zaixiang Zheng, Yifan Deng, Dongyu Xue, Yi Zhou, Fei Ye, and Quanquan Gu. Structure-informed language models are protein designers. bioRxiv, pages 2023–02, 2023.

39. Jeffrey M Spencer and Xiaoliu Zhang. Deep mutational scanning of s. pyogenes cas9 reveals important functional domains. Scientific reports, 7(1):16836, 2017.

40. Li Xu, Yakun Liu, and Renzhi Han. Beat: a python program to quantify base editing from sanger sequencing. The CRISPR journal, 2(4):223–229, 2019.

41. David Conant, Tim Hsiau, Nicholas Rossi, Jennifer Oki, Travis Maures, Kelsey Waite, Joyce Yang, Sahil Joshi, Reed Kelso, Kevin Holden, et al. Inference of crispr edits from sanger trace data. The CRISPR journal, 5(1):123–130, 2022.

42. Bowen Jing, Stephan Eismann, Patricia Suriana, Raphael John Lamarre Townshend, and Ron Dror. Learning from protein structure with geometric vector perceptrons. In International Conference on Learning Representations, 2020.

